# Synthetic metabolic pathways for conversion of CO_2_ into secreted short-to medium-chain hydrocarbons using cyanobacteria

**DOI:** 10.1101/2021.07.08.451587

**Authors:** Ian S. Yunus, Josefine Anfelt, Elton P. Hudson, Patrik R. Jones

**Affiliations:** Department of Life Sciences, Imperial College London, SW7 2AZ, London, United Kingdom; KTH - Royal Institute of Technology, Division of Proteomics and Nanobiotechnology, Science for Life Laboratory, Stockholm SE-171 21, Sweden

**Keywords:** Propane, Propene, gasoline, renewable, biofuels, CRISPRi, dCas9, cyanobacteria, metabolic engineering

## Abstract

The objective of this study was to implement direct sunlight-driven conversion of CO_2_ into a naturally excreted ready-to-use fuel. We engineered four different synthetic metabolic modules for biosynthesis of short-to medium-chain length hydrocarbons in the model cyanobacterium *Synechocystis* sp. PCC 6803. In module 1, the combination of a truncated clostridial n-butanol pathway with over-expression of the native cyanobacterial aldehyde deformylating oxygenase resulted in small quantities of propane when cultured under closed conditions. Direct conversion of CO_2_ into propane was only observed in strains with CRISPRi-mediated repression of three native putative aldehyde reductases. In module 2, three different pathways towards pentane were evaluated based on the polyunsaturated fatty acid linoleic acid as an intermediate. Through combinatorial evaluation of bioreaction ingredients it was concluded that linoleic acid undergoes a spontaneous non-enzymatic reaction to yield pentane and hexanal. When *Synechocystis* was added to the bioreaction, hexanal was converted into 1-hexanol, but there was no further stimulation of pentane biosynthesis. For modules 3 and 4, several different acyl-ACP thioesterases were evaluated in combination with two different decarboxylases. Small quantities of 1-heptene and 1-nonene were observed in strains expressing the desaturase-like enzyme UndB from *Pseudomonas mendocina* in combination with C8-C10 preferring thioestersaes. When UndB instead was combined with a C12-specific ‘*Uc*FatB1 thioesterase, this resulted in ten-fold increase of alkene biosynthesis. When UndB was replaced with the light-dependent FAP decarboxylase, both undecane and tridecane accumulated, albeit with a 10-fold drop in productivity. Optimization of the RBS, promoter and gene order in these synthetic operons resulted in 1-alkene bioproductivity of 230 mg/L after 10 d with 15% carbon partitioning. In conclusion, the direct bioconversion of CO_2_ into secreted and ready-to-use hydrocarbon fuel was accomplished and optimal results were obtained with UndB and a C12 chain-length specific thioesterase.

**Highlights:**

1. Multiple repression of endogenous aldehyde reductases/dehydrogenases by CRISPRi enabled propane biosynthesis
2. Biosynthesis of short-medium chain hydrocarbons (C7-C11) in a cyanobacterium was demonstrated for the first time
3. The final enzymes of the hydrocarbon pathways influenced both productivity and product profile
4. All volatile products were naturally secreted and accumulated outside of the cell

## 1. Introduction

Industrial (white) biotechnology, defined as the use of engineered biology with or without cells for the provision of fuels, chemicals, and materials, provides an opportunity to reduce our dependency on unsustainable sourcing from fossil fuels or tropical plant agriculture. However, in many cases, this is challenging due to high manufacturing costs for biotechnology-based products and incomplete financial accounting in both fossil fuel and tropical plant agriculture-based systems, the latter resulting in unfair competition. For example, neither producers nor end-users pay for all the indirect costs associated with manufacturing and use of current products (Perin and Jones, 2019). Instead, all of society pay those costs. Hence, such indirect costs are not reflected in the price paid by end-users. This contributes towards keeping unit costs low and the threshold for competition high, therefore making it difficult to commercialize new biotechnology-based products. Obviously, this is only relevant if industrial biotechnology is more sustainable (Ögmundarson et al., 2020), which can be difficult to properly assess without large-scale implementation and subsequent optimization of practice.

Consequently, biotechnology-based business and research is increasingly shifting toward higher value, lower quantity products, where the chance of successful commercialization is greater. This is important and can provide excellent opportunities to increase the industry base and its experience and techniques for optimized operation. Whilst so doing, it remains important to not forget the earlier ambition to develop and improve biotechnology also for higher quantity products, such as fuel, since this is where the greatest need for impact remains.

Part of the reason why biotechnological chemicals and fuel manufacturing is financially and energetically costly relate to downstream processing such as cell harvesting, extraction, chemical processing incl. distillation, for recovery of sufficiently purified products (Fasaei et al., 2018; Kim et al., 2013). One approach to mitigate such costs is to pursue fuels that (1) naturally secrete from the cell, (2) easily can be separated from the aqueous media, and (3) are ready-to-use without further processing. Several shorter chain-length alkanes, like C3 propane, are volatile and there is already an existing market. Several different synthetic pathways for short-chain length alkanes have been implemented in heterotrophic microorganisms. For example, synthetic pathways enabling the biosynthesis of C3 to C13 alkanes in heterotrophic microorganisms have been reported (Kallio et al., 2014; Schirmer et al., 2010; Sheppard et al., 2016; Zhu et al., 2017).

Looking even further into the future, industrial biotechnology should ideally not compete for resources (e.g., water, land, nutrients) with agriculture, especially given that with increasing climate change it may become limited, whilst the population continues to grow. The commercial use of engineered photosynthetic microorganisms (eukaryotic algae and cyanobacteria) for biotechnological manufacturing is today very limited. Engineered photosynthetic biosynthesis of alkanes with a chain-length of C11 and above have been described (Yunus et al., 2018), however, none of the products were found to be excreted (C13-C17), or their accumulation was negligible (C11). Photosynthetic biosynthesis of C3 propane has also recently been reported, using a fatty acid photodecarboxylase (FAP) (Amer et al., 2020).

With an eye towards eventually enabling photosynthetic conversion of CO_2_ directly into naturally secreted and ready-to-use fuels, we here report the initial development of four metabolic systems (Fig. 1) for short-to-medium chain-length hydrocarbon biosynthesis in the model cyanobacterium *Synechocystis* sp. PCC 6803.

**Figure 1.**
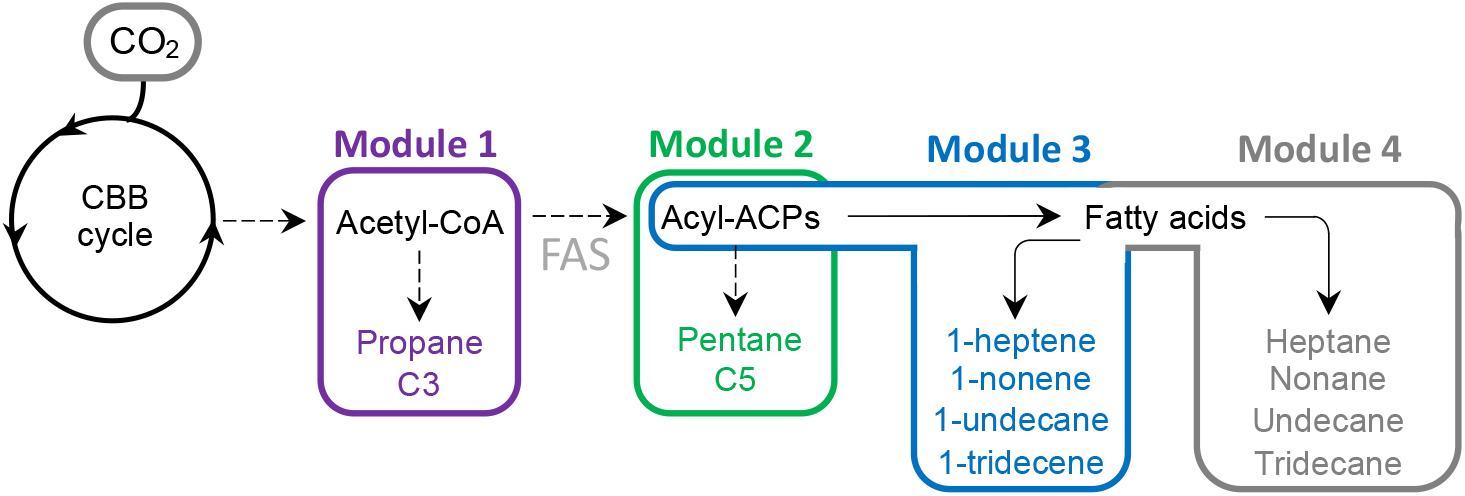
Graphic overview of chemical product targets and pathways thereto.

## 2. Materials and methods

### 2.1. Plasmid propagation

*E. coli* DH5α (Thermo Fisher Scientific) was used for propragation of all plasmids used in the study. The *E. coli* DH5α strains were routinely cultivated in lysogeny broth (LB) medium (LB Broth, Sigma Aldrich), 37°C, 180 rpm, and supplemented with appropriate antibiotic(s) (final concentration: spectinomycin 50 μg/mL, carbenicillin 100 μg/mL, chloramphenicol 37 μg/mL, and erythromycin 200 μg/mL). Plasmids were constructed using a modular plasmid assembly method, namely Biopart Assembly Standard Idempotent Cloning (BASIC) as described previously (Storch et al., 2015; Yunus et al., 2020, 2018; Yunus and Jones, 2018) unless stated otherwise. All amino acid sequences of enzymes used in this study are listed in Supplementary Table S1.

### 2.2 *Cultivation condition for Synechocystis* sp. PCC 6803

All *Synechocystis* sp. PCC 6803 strains were cultivated in BG11 medium without cobalt (hereafter BG11-Co) as cobalt was used as an inducer in some of the engineered strains. Prior to production experiment, liquid preculture was grown in a 6-well plate in BG11-Co containing appropriate antibiotic(s) (spectinomycin 25 μg/mL, kanamycin 50 μg/mL, chloramphenicol 37 μg/mL, and erythromycin 20 μg/mL) at 30 °C, 180 rpm (Unimax 1010 shaker, Heidolph Instruments), 60 μE (warm-white LEDs) in AlgaeTron AG 230 (Photon Systems Instruments (PSI), Czech Republic). The AlgaeTron AG 230 was equipped with CO_2_ sensor and controller (Ecotechnics, UK) to control and monitor the concentration of CO_2_ (1% (v/v) inside the growth chamber. Liquid preculture (OD_730_ 3-4) was transferred to a 100-mL Erlenmeyer flask for production experiments and the OD_730_ was adjusted to 0.2 by adding BG11-Co medium to a final volume of 25 mL with antibiotic(s). Production experiments were conducted under the same condition unless stated otherwise.

For propane production experiments, liquid preculture (OD_730_ 3-4) was transferred to a 100-mL Erlenmeyer flask (E-flask) or a 25 cm^2^ Tissue Culture flask (T-flask) with a plug seal (Thermo Fisher Scientific), and the OD_730_ was adjusted to 0.2 by adding BG11-Co medium to a final volume of 25 mL with antibiotic(s). The cultures were induced on day two with 1 mM isopropyl β-D-1-thiogalactopyranoside (IPTG) and 1 mg/L anhydrotetracycline (aTc), cultivated for four days, sampled for n-butanol and butyraldehyde analysis, and were transferred (7.5 mL) to serum bottles (size: 75 mL). The serum bottles were sealed with gastight silver aluminium rubber stoppers and the cultures were then allowed to grow for 48 hrs prior to propane measurement from the headspace using GC-FID.

For pentane production experiments, liquid cultures and control samples in 100-mL Erlenmeyer flasks were cultivated for six days before 7.5 mL of the liquid culture was transferred to serum bottles. Linoleic acid (180 mg/L, dissolved in 100% ethanol) was added to the serum bottles unless otherwise stated. The closed vessels were cultivated for two more days with and without isopropyl myristate overlay (10% v/v). Hexanal and 1-hexanol were monitored from the solvent overlay and analyzed using GC-MS, whilst pentane was monitored using GC-FID from the headspace of samples containing no overlay.

For production experiments of terminal alkenes, liquid cultures in 100-mL Erlenmeyer flasks were induced with cobalt (5 μM) and overlaid with 10% (v/v) hexadecane on day two. The terminal alkenes were sampled from the solvent overlay on day 10 and analyzed using GC-MS.

For production experiments of fatty alkanes, liquid cultures in 100-mL Erlenmeyer flasks were cultivated under continous illumination of warm white LEDs (60 μE) and blue light LEDs (100 – 150 μE). Blue light was used to activate the fatty acid photodecarboxylase (FAP) enzyme. On day two, 625 nM cobalt was added to drive the expression of *‘Cp*FatB1.4 thioesterase under Pcoa, a cobalt-inducible promoter, and the hexadecane solvent overlay (10% v/v) was added. In some cases, multiple inductions were also performed. Alkanes were anaylzed from the solvent overlay on day 10 using GC-MS.

### 2.3 T*ransformation methods for Synechocystis* sp. PCC 6803

Chromosomal integration and transformation of RSF1010 plasmids were performed as described previously (Sattayawat et al., 2020; Yunus et al., 2020, 2018; Yunus and Jones, 2018). All strains generated in this study are listed in Supplementary Table S2.

### 2.4. Product analysis

For analysis of fatty alkanes, terminal alkenes, 1-hexanol, or hexanal from the solvent overlay, 100 μL of solvent overlay was sampled and transferred into a vial insert (5183-2085, Agilent) in a GC vial. Sample (1 μL) was analyzed using Hewlett Packard (HP) 6890 Series Gas Chromatograph (GC) equipped with a 5973 Mass Selective Detector and a DB-WAXetr column (Agilent, 122-7332). The injector temperature was set at 246 °C and helium was used as the carrier gas (1.5 mL/min). The oven temperature was initially held at 70°C for 30 s, followed by a ramp at 30°C.min^-1^ to 250°C and hold for 1 min. Serial dilutions of hydrocarbon, 1-hexanol, and hexanal standard were also prepared to quantify the concentration of fatty alkanes, terminal alkenes, or 1-hexanol in the liquid culture.

For analysis of propane and pentane, 500 μL of headspace sample was taken and analyzed using a 8860 Gas Chromatograph (GC) System equipped with a Flame Ionization Detector (FID) and a GS-GasPro UST 1446134 column (Agilent, 113-4332). The inlet temperature was set at 250 °C (8.7 psi). Flow rate was maintained at 22.143 ml/min, with 0.743 ml/min purge flow. For propane and pentane analysis, oven was set at 90 °C and 120 °C, respectively, and hold for 5 min. Propane and pentane standard was purchased from BOC gases (United Kingdom) and Sigma Aldrich, respectively.

For analysis of 1-butanol, 1 mL of liquid culture was harvested by centrifugation at 17,000 x *g* for 20 min. The supernatant (750 μL) was transferred into a HPLC vial. Sample (100 μL) was analyzed using an Agilent 1200 Series HPLC instrument equipped with an Aminex HPX-87H column (Bio-Rad). Water containing 5 mM sulphuric acid was used as the mobile phase. Flow rate and column temperature was set at 0.6 ml/min and 60 °C, respectively. A serial dilution of 1-butanol was prepared to quantify the concentration of 1-butanol in the liquid culture.

## 3. Results

### 3.1 Module 1 – CRISPRi-mediated biosynthesis of propane using Ado

Three different synthetic metabolic routes for bioproduction of propane have been reported (Amer et al., 2020; Kallio et al., 2014; Menon et al., 2015; Sheppard et al., 2016), differentiated by the central carbon metabolic precursor (valine, acyl-ACP or acyl-CoA) and/or the type of enzyme used in the last catalytic step (Ado or FAP). The first pathway requires an aldehyde deformylating oxygenase (Ado) and utilizes an aldehyde (e.g., butyraldehyde) as the substrate (Kallio et al., 2014; Menon et al., 2015; Sheppard et al., 2016). The second requires a blue-light dependent fatty acid photodecarboxylase (FAP) (Sorigué et al., 2017) that catalyzes the decarboxylation of carboxylic acids (e.g. butyric acid or isobutyric acid) to propane (Amer et al., 2020). Recently it was proposed that the native substrate of FAP is acyl-CoA (Li et al., 2020). Although the FAP-dependent pathway proceeds in a simpler manner (i.e., requires fewer catalytic steps) and has been implemented for propane production in cyanobacteria (Amer et al., 2020), the improved FAP_G462V_ enzyme suffers from photoinactivation at higher light intensities (Amer et al., 2020) which likely limits FAP-dependent propane production in cyanobacteria. To avoid this limitation and evaluate an alternative route, we set out to develop synthetic pathways for propane production in *Synechocystis* sp. PCC 6803 (hereafter *Synechocystis*) that instead utilize Ado.

The Ado-dependent pathway to propane requires butyraldehyde as the penultimate precursor. We therefore constructed a butyraldehyde biosynthesis pathway (Fig. 2A) based on the modified Clostridial fermentation pathway for *n*-butanol biosynthesis previously engineered in *E. coli* (Atsumi et al., 2008; Nielsen et al., 2009), yeast (Krivoruchko et al., 2013; Steen et al., 2008), and cyanobacteria (Anfelt et al., 2015; Lan et al., 2013; Lan and Liao, 2012, 2011). The initial step involved over-expression of *xfpK* (phosphoketolase, *Bifidobacterium breve*, UniprotKB ID: D6PAH1) to increase the acetyl-CoA pool concentration as shown previously (Anfelt et al., 2015). Next, the endogenous *phaA* gene (acetoacetyl-CoA thiolase, encoded by *slr1993*) was knocked out to avoid the thermodynamically unfavorable condensation reaction of two molecules of acetyl-CoA to acetoacetyl-CoA by native PhaA (Lan and Liao, 2012). Acetyl-CoA is naturally converted into malonyl-CoA by AccABCD (acetyl-CoA carboxylase). Condensation of malonyl-CoA and acetyl-CoA into acetoacetyl-CoA was achieved by over-expression of NphT7 (acetoacetyl-CoA synthase, *Streptomyces* sp. Strain CL190, UniprotKB ID: D7URV0). This condensation reaction is irreversible and was reported to stimulate 1-butanol synthesis in cyanobacteria (Lan and Liao, 2012). Next, acetoacetyl-CoA was converted to butyraldehyde by native PhaB (native acetoacetyl-CoA reductase, *slr1994*) and heterologously expressed PhaJ (enoyl-CoA hydratase, *Aeromonas caviae*, UniProtKB ID: O32472), Ter (trans-enoyl-CoA reductase, UniProtKB ID: Q73Q47), and PduP (O2-tolerant aldehyde dehydrogenase, *Salmonella enterica*, UniProtKB ID: V2D4V9). Finally, butyraldehyde was converted into propane by Ado (aldehyde deformylating oxygenase) obtained from *Nostoc punctiforme* (UniProtKB ID: B2J1M1) or *Prochlorococcus marinus* (UniProtKB ID: Q7V6D4). To minimize the conversion of butyraldehyde to 1-butanol, several endogenous *ahr* genes (aldehyde reductase/dehydrogenase, encoded *slr0942, sll0990, slr0091*, and *slr1192)* were repressed using CRISPRi. Addditionally, native *phaEC* genes (poly(3-hydroxyalkanoate) polymerase subunit E and C, encoded by *slr1829* and *slr1830*, respectively) were also knocked out to minimize the biosynthesis of polyhydroxybutyrate (PHB).

**Figure 2.**
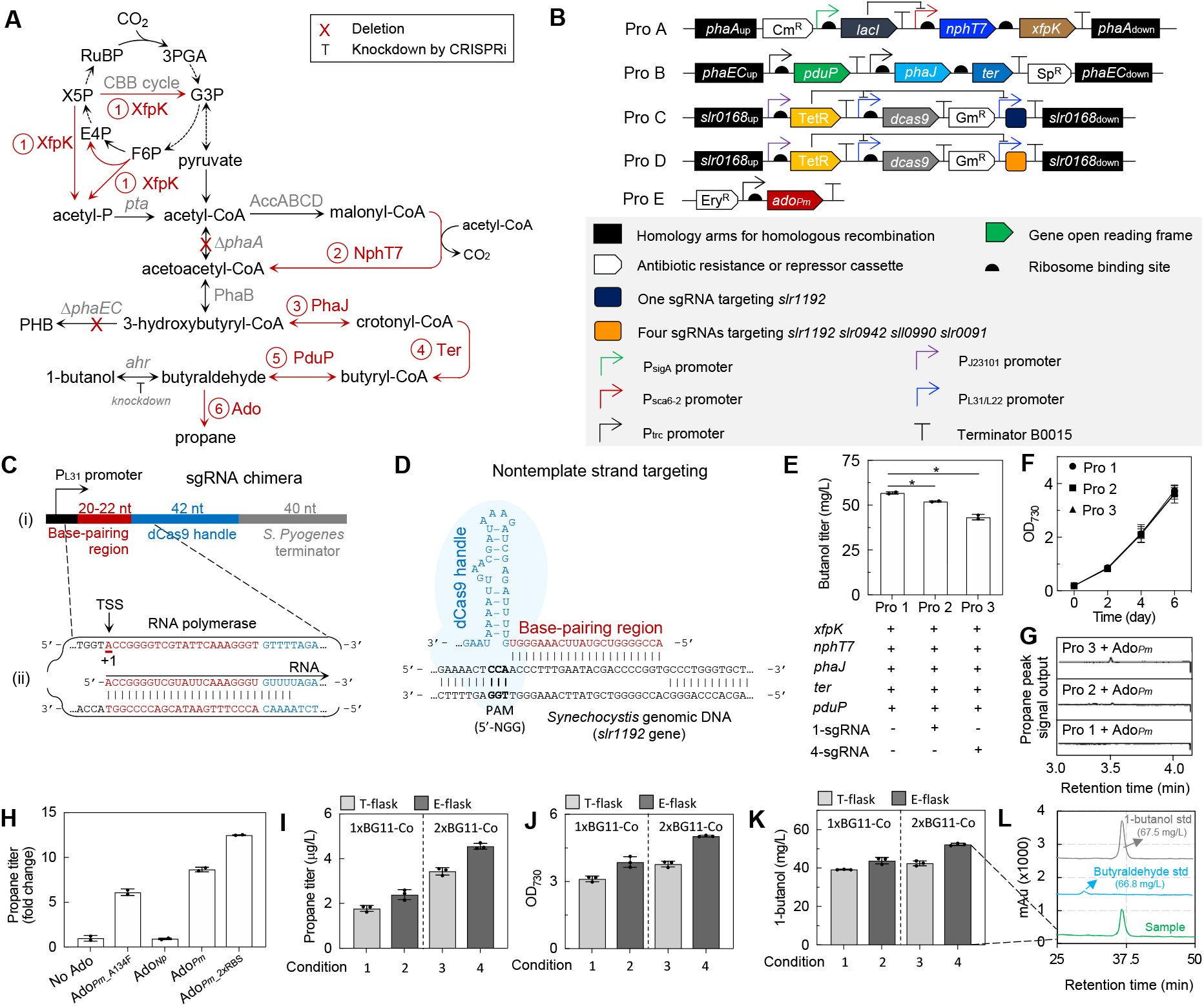
Strategies for ADO-dependent propane production in *Synechocystis* sp. PCC 6803. (A) Synthetic metabolic pathways of ADO-dependent propane biosynthesis. Endogenous genes are written in gray, whilst heterologous genes are numbered and written in red. (B) Schematic diagram of plasmids used to generate strains shown in Fig. 2E-G. (C) An example of sgRNA chimera design for targeting *slr1192* gene: (i) sgRNA consists of a P_L31_ promoter (black), a 20 nt base-pairing region for specific DNA binding (red), a 42 nt encoding hairpin structure for dCas9 binding (blue), and a 40 nt transcription terminator derived from *S. pyogenes* (gray), (ii) P_L31_ promoter requires an A nucleotide as the transcription start site (TSS or +1) at the 5’ end of the transcript for effective transcription. This TSS was taken into consideration when searching for a target in the nontemplate strand. (D) The pattern 5’-A-N_(19-24)_-NGG-3’ from the reverse strand in *slr1192* gene was used to find the sequence for base-pairing region. The reverse-complement sequence of A-N_(19-24)_ was used as the base-pairing region of the sgRNA. The transcribed sequence of base-pairing region in Fig. 2C (ii) binds to the nontemplate strand in Fig. 2D. (E) Butanol titer obtained from Strain Pro 1, 2, and 3. (F) Growth profile of strains shown in Fig. 2E. (G) Propane peak intensity upon overexpression of *ado_PM_* in Strain Pro 1, 2, and 3. (H) Selection of different Ado variants by feeding 160 mg/L butyraldehyde. (I) Propane titer, (J) cell density, and (K) 1-butanol titer of Pro 3 + Ado_*Pm_2xRBS*_ when cultivated in T-flask and E-flask without exogenous feeding of butyraldehyde. (L) 1-butanol and butyraldehyde analysis of Sample 4 in Fig. 2K by HPLC. Data were obtained from two-three biological replicates and error bars represent standard deviation.

The above described engineering was realized in *Synechocystis* using four integrative plasmids targeting the *phaA, phaEC*, and *slr0168* sites (Fig. 2B, Pro A-D) and one self-replicating RSF1010 plasmid (Fig. 2B, Pro E). Plasmid Pro A was used to overexpress both *xfpk* and *nphT7* under P_sca6-2_, an IPTG-inducible promoter (Albers et al., 2015), while simultaneously knocking out the *phaA* gene. Likewise, plasmid Pro B was used to overexpress *pduP, phaJ, and ter* genes under a P_trc_ promoter, and integrated into the *phaEC* site to eliminate PHB biosynthesis. To lower aldehyde reduction by native reductase and dehydrogenases, we used a CRISPRi method described previously (Yao et al., 2016). Genes encoding dCas9 (a catalytically dead Cas9) and the single guide RNA (sgRNA) were integrated into the *slr0168* site under the control of an anhydrous tetracycline (aTc)-inducible P_L31/L22_ promoter (Huang and Lindblad, 2013). The sgRNA(s) targeting one (*slr1192*; Fig. 2C,D) or four native annotated aldehyde reductases/dehydrogenases (*slr0942, sll0990, slr0091*, and *slr1192*) were designed (Supplementary Note S1). The sgRNA constructs consisted of a 20-22 nt base-pairing region, a 42 nt dCas9 handle, and a 40 nt Rho-independent transcription terminator (Fig. 2C panel (i)) as described previously (Larson et al., 2013; Qi et al., 2013). All sgRNAs were designed to target the nontemplate strand, as this was previously reported more effective for repression in *E. coli* (Bikard et al., 2013; Qi et al., 2013), and the base-pairing region (Fig. 2C panel (ii), highlighted in gray) was designed as the reverse-complementary sequence of the target DNA in the nontemplate strand (Fig. 2D, highlighted in gray). Finally, *ado_Pm_* from *P. marinus* was cloned into a broad-host range self-replicating RSF1010 plasmid under a constitutive Ptrc promoter (Fig. 2B, Pro E).

Transformed strains were initially cultivated in Erlenmeyer flasks (E-flask) and thereafter transferred into serum bottles sealed with gas-tight PTFE caps prior to propane measurement. We found that both strains expressing 1-sgRNA (Strain Pro 2; Fig. 2E) or 4-sgRNAs (Strain Pro 3; Fig. 2E) produced significantly less butanol compared to the control strain (Strain Pro 1; Fig. 2E), although they all grew similarly (Fig. 2F). Upon overexpression of *ado_Pm_*, the strain with 4-sgRNA showed the highest accumulation of propane (Fig. 2G), whilst no propane was observed from the Pro 1 + Ado_*Pm*_ strain. While it is known that *slr1192* gene is involved in biosynthesis of n-butanol (Liu et al., 2019) and isobutanol (Miao et al., 2017), little is known about the function of slr0942, sll0990, and slr0091 in butyraldehyde/n-butanol biosynthesis pathway. Our study suggests that the gene product of *slr0942, sll0990*, and *slr0091*, that is involved in production of volatile alcohols during lipid peroxidation (Hintzpeter et al., 2015) or in ethanol production (Gao et al., 2012) and conversion of apocarotenals and alkanals into the corresponding acids (Trautmann et al., 2013), might also play roles in butanol production and that the repression of these genes facilitated the accumulation of butyraldehyde in the liquid culture.

Encouraged by this finding, we therefore continued to search for better Ado candidates by overexpressing several variants of Ado(s), including the Ado_A134F_ mutant from *P. marinus* which was found to have high specificity for short-to medium-chain aldehydes (Khara et al., 2013). Cultures expressing the different Ado candidates were spiked with approximately 160 mg/L butyraldehyde following transfer to serum bottles. In contrast to earlier reports (Khara et al., 2013), strains expressing wild-type Ado accumulated a higher propane titer than those expressing the AdoA134F variant (Ado_*Pm*_ vs Ado_A134F_; Figure 2H). By placing one additional ribosome-binding site (RBS) upstream of the *ado_Pm_* gene to enhance its protein expression, we further improved the propane production by nearly 1.3-fold compared to the strain with only one RBS (Ado_*Pm*_2xRBS_ vs Ado_*Pm*_, Figure 2H, Supplementary Fig. S1). Whether or not these differences were caused by changes in specific or total activities remains unknown given that the accumulation of Ado was not quantified in any strain.

After optimizing the choice of Ado, we evaluated the effect of environmental conditions on 1-butanol/butyraldehyde production. Studies by Miao and colleagues (Miao et al., 2017) demonstrated that the concentration of volatile alcohols such as isobutanol decreases over time when cultivated in an open system, such as Erlenmeyer flasks (E-flasks). When incubated in a closed system, such as Tissue Culture flasks (T-flasks) with plug seal caps, the concentration of isobutanol remained relatively constant after 9 days of incubation. We hypothesized that since butyraldehyde has an even higher vapor pressure than isobutanol, it might also have been lost during the six-day cultivation in E-flasks. Therefore, to minimize the loss of butyraldehyde during a six-day cultivation in the E-flasks, we evaluated the production system also in T-flask immediately before transfer to serum bottles. Curiously, Figure 2I shows that the propane titer produced in E-flasks was higher than that from T-flask. This might be attributed to the fact that the OD_730_ of cells cultivated in E-flask was higher (Fig. 2J). Indeed, when we cultivated the strains in 2xBG11-Co, the cells grow better (Fig. 2J) and produced even higher propane titer (up to 4.5 ug/L) (Fig. 2I). When we measured the accumulated butyraldehyde in the Erlenmeyer flasks, however, we found that a butyraldehyde peak could not be detected from any samples, possibly due to being lower than its the detection limit. Although the propane titer achieved in this study (4.5 μg/L) was substantially lower than the titer achieved from the FAP-dependent pathway, 11 mg/L/day under open continuous conditions (Amer et al., 2020), we demonstrated that CRISPRi-based repression of native aldehyde reductases improves propane production. Similarly to what was reported earlier for medium-to-long chain-length hydrocarbons (Yunus et al., 2018), synthetic pathways utilizing the algal FAP enzyme were substantially superior to those utilizing the cyanobacterial ADO enzyme.

### 3.2 Module 2 – Biosynthesis of pentane via a lipoxygenase pathway

There are at least two naturally occurring and one synthetic pathways that have been reported for pentane production: (1) the long-known pentane evolution of tissue lipids from ω6 fatty acids during lipid peroxidation (Horvat et al., 1964), (2) a direct lipoxygenase-mediated conversion of linoleic acid (C18:2(9,12)) to pentane in soybean plants (*Glycine max*) (Sanders et al., 1975) which has been reconstructed in *Yarrowia lipolytica* (Blazeck et al., 2013), and (3) a synthetic hexanal-based pentane biosynthesis using Ado and a modified clostridial n-butanol pathway (Sheppard et al., 2016). In this study, we explored the possibility of producing pentane from linoleic acid in *Synechocystis* sp. PCC 6803.

In plants, linoleic acid released from membrane lipids would normally undergo oxidation in response to biotic or abiotic stress, which results in the sequential transformation of polyunsaturated fatty acids into signalling and defense molecules, such as volatile C6 or C9 aldehydes, alcohols, or esters – the so-called green leaf volatiles (GLVs) (Feussner and Wasternack, 2002). Cyanobacteria can be engineered to accumulate unsaturated free fatty acids by modification of the fatty acid biosynthesis pathway and overexpression of desaturase enzymes (Chen et al., 2014; Santos-Merino et al., 2018; Yoshino et al., 2017). The best strain can accumulate up to 30% unsaturated fatty acids (Santos-Merino et al., 2018), making cyanobacteria a promising host for pentane production.

To minimize the recycling of linoleic acid back to the fatty acid biosynthesis pathway, we first knocked out the acyl-ACP synthetase gene (*aas*) which resulted in the secretion of free fatty acids to the liquid medium (Kaczmarzyk and Fulda, 2010) (Fig. 3A). Next, we designed experiments to investigate the conversion of linoleic acid to pentane via three different routes: (1) lipoxygenation of linoleic acid, with the assumption that *Synechocystis* also generates GLVs, and that some of the resulting C6 or C9 aldehydes are converted into corresponding hydrocarbons by an aldehyde deformylating oxygenase (Fig. 3B (i)), (2) direct lipoxygenation of linoleic acid by a soybean lipoxygenase (*Gm*Lox1) without any aldehyde or alcohol by-products (Fig. 3B (ii)), and (3) non-enzymatic reaction(s) of linoleic acid to pentane and other by-products (Donovan and Menzel, 1978) (Fig. 3B (iii)).

**Figure 3.**
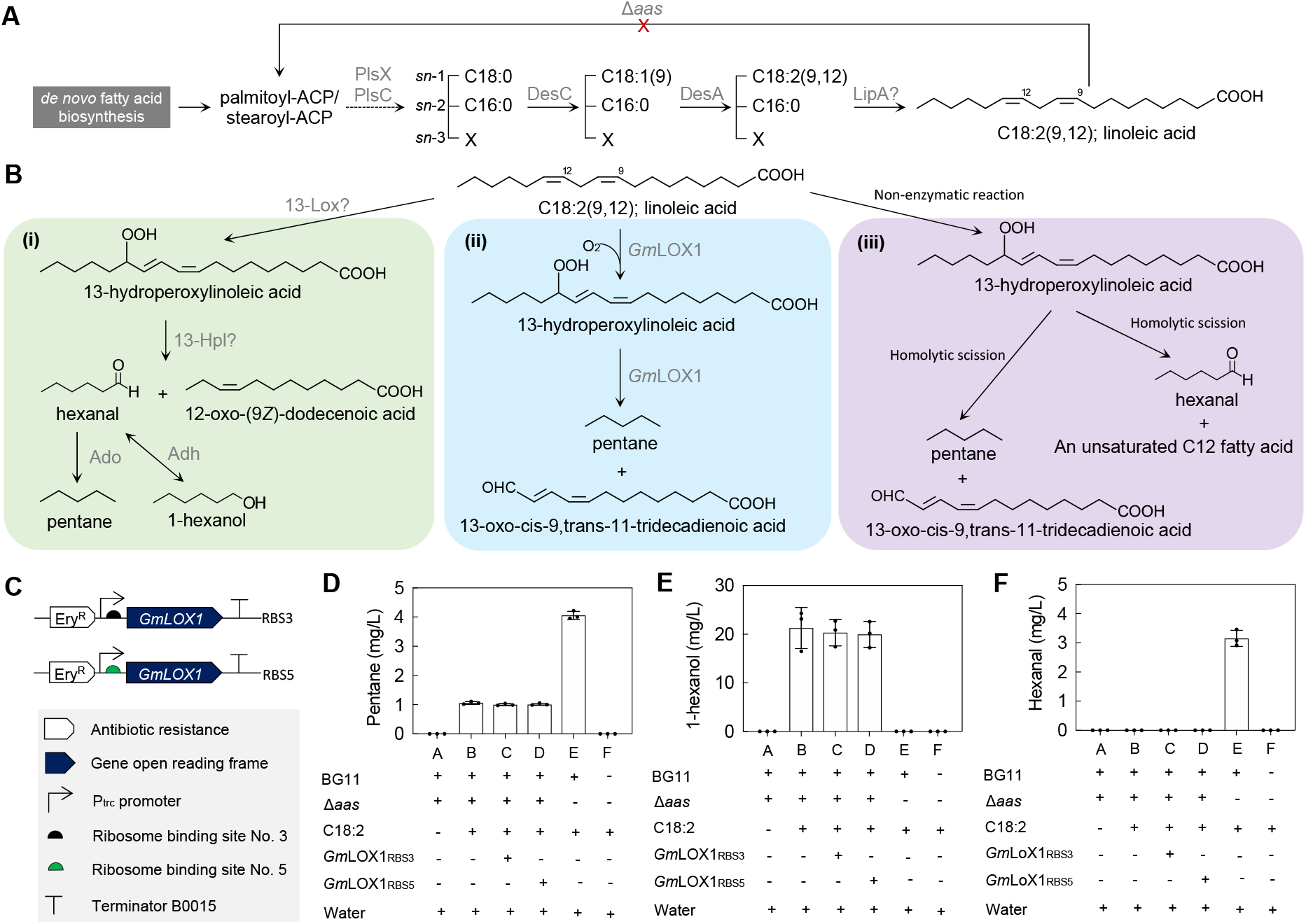
Non-enzymatic conversion of exogenous linoleic acid into pentane. (A) Linoleic acid biosynthesis from *de novo* fatty acid biosynthesis pathway. (B) Three alternative routes for pentane biosynthesis: (i) lipoxygenase pathway, (ii) soybean lipoxygenase pathway, (iii) non-enzymatic reaction. (C) Schematic diagram of plasmid for *GmLOX1* overexpression. (D) Pentane, (E) 1-hexanol, and (F) hexanal titer obtained from different sample treatments. Liquid cultures were grown in E-flasks for 6 days and transferred into serum bottles. For pentane quantification, headspace was sampled and analyzed using GC-FID. For 1-hexanol or hexanal quantification, liquid cultures were overlaid with isopropyl myristate. Isopropyl myristate solvent overlay was sampled and analyzed using GC-MS. Samples were taken on day 8. Data were obtained from three biological replicates and error bars represent standard deviation. PlsX; putative phosphate acyltransferase (*slr1510*), PlsC; putative 1-acyl-sn-glycerol-3-phosphate acyltransferase (*sll1848*), DesC; Δ9 acyl lipid desaturase (*sll0541*), DesA; Δ12 acyl lipid desaturase (*slr1350*), LipA; putative lipase (*sll1969*), 13-Lox; putative 13-lipoxygenase, 13-Hpl; putative 13-hydroperoxide lyase, *Gm*LOX1; 13-lipoxygenase from *Glycine max* (Accession No. P08170)

We grew Δ*aas* liquid cultures in E-flasks for 6 days before transfer into serum bottles. The closed vessels were incubated for 48 hrs with and without isopropyl myristate solvent overlay. Headspace samples from serum bottles without solvent overlay were taken for pentane analysis, whilst 1-hexanol and hexanal were monitored by sampling the overlay fraction. The results (treatment A, Fig. 3D-F) indicate that pentane (Fig. 3D), 1-hexanol (Fig. 3E), or hexanal (Fig. 3F) were not observed, neither from the headspace nor the solvent overlay (Supplementary Fig. S2). We hypothesized that the level of naturally generated linoleic acid was too low to generate detectable pentane, hexanal, or hexanol. To increase the concentration of linoleic acid, we then repeated the same closed cultures and spiked each one of them with approximately 180 mg/L of linoleic acid. Surprisingly, the highest pentane titer was obtained in treatment E, which only contains BG11 and linoleic acid without *Synechocystis* cells (Fig. 3D). Hexanal (Fig. 3F), but not 1-hexanol (Fig. 3E), was also observed from this sample, indicating that in the absence of *Synechocystis* cells, most linoleic acid undergoes a non-enzymatic reaction to yield pentane and hexanal. In the presence of *Synechocystis* (treatment B), no hexanal was observed (Fig. 3F) and around 20 mg/L 1-hexanol accumulated instead. Curiously, upon overexpressing *GmLOX1*, which was expected to streamline the conversion of linoleic acid to pentane, we did not see any improvement of pentane production from treatments C and D in comparison to B (Fig. 3D). Thus, further work is needed to implement a direct route from CO_2_ to pentane in cyanobacteria, either by enhancing precursor accumulation and/or by identifying a LOX candidate which works effectively. Such work could also benefit from combination with the 1- and 4-sgRNA constructs to minimize the loss of ADO-precursors into corresponding alcohols. Alternatively, the biosynthesis of hexanoic acid in combination with FAP is another possibility to explore.

### 3.3 Module 3 – Biosynthesis of C7-C13 terminal alkenes with UndB

The two previous modules described metabolic pathways for the production of propane and pentane. Both pathways share similarities, in that some of them are Ado-dependent and repression of multiple aldehyde reductases were either found to or should most likely maximize the hydrocarbon accumulation. In an earlier study with longer chain-length fatty acid substrate we found that the desaturase-like enzyme UndB from *Pseudomonas mendocina* (Rui et al., 2015) was highly effective in synthesizing terminal alkenes in *Synechocystis* sp. PCC 6803 (Yunus et al., 2018).

To understand the utility of UndB for shorter chain-length products we fed a range of free fatty acids (C4:0-C18:0) to a culture of *Synechocystis* over-expressing UndB (Fig. 4A-B). The results suggest that UndB is active on fatty acids with a broad range of chain-lengths. The highest activity was observed with dodecanoic acid (C12:0) (Fig. 4C), in line with previous studies (Rui et al., 2015; Yunus et al., 2018). Interestingly, and which to our understanding has not yet been reported elsewhere, we found that UndB was also active on butyric acid resulting in the synthesis of propene, although very weakly (Supplementary Fig. S3).

**Figure 4.**
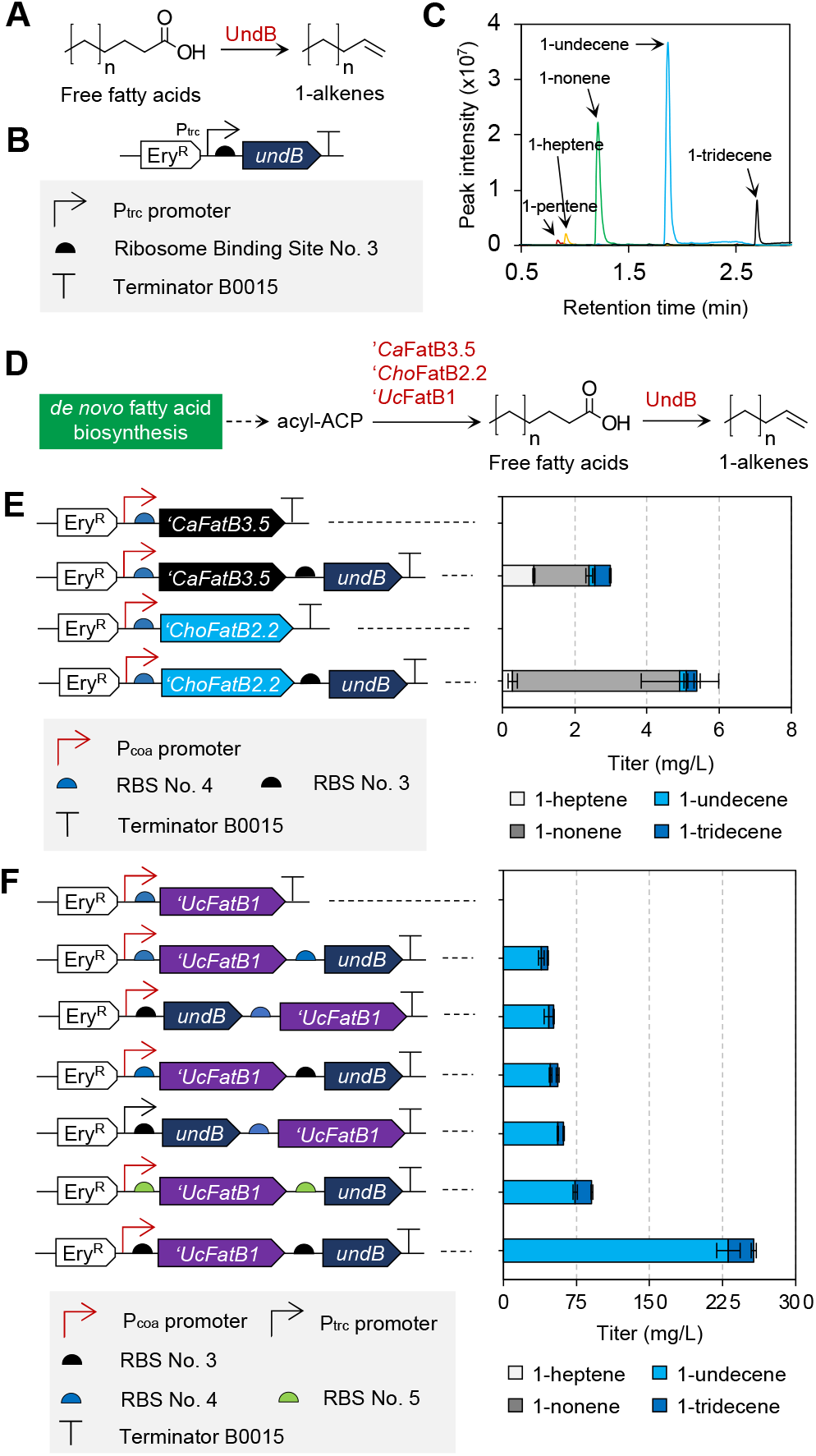
Production of 1-alkenes in *Synechocystis*. (A) Conversion of free fatty acids to 1-alkenes by feeding different chain length of free fatty acids to a strain expressing UndB (B). (C) Chromatogram obtained from GC-MS. (D) Synthetic pathway of 1-alkenes biosynthesis. Acyl-ACPs were converted into their corresponding free fatty acids by a thioesterase followed by conversion of free fatty acids into 1-alkenes by UndB. (E, F) Production of 1-heptene, 1-undecene, 1-nonene from strains expressing different thioesterases and UndB. For feeding experiments (Fig. 4C), liquid cultures were spiked with 100 mg/L free fatty acid and overlaid with 10% (v/v) hexadecane on day two. Samples were collected from the solvent overlay on day ten and analyzed by GC-MS. For production experiments (Fig. 4E,F), liquid cultures were induced with 5 μM cobalt on day two, overlaid with 10% (v/v) hexadecane, and cultivated until day ten before analysis of the solvent overlay. The growth profile of some strains shown in Fig. 4F is presented in Supplementary Fig. S4. Data were obtained from three biological replicates and error bars represent standard deviation.

To realize the biosynthesis of short and medium chain-length 1-alkenes in *Synechocystis*, UndB was co-expressed together with three different thioesterases which previously have been reported to display specificity toward short and medium chain-length acyl-ACP (Fig. 4D). This includes ‘*Ca*FatB3.5, a thioesterase that when truncated in the optimal position is capable of generating a broad range of fatty acids in cyanobacteria (Yunus et al., 2021) and was previously used for bioproduction of 1-octanol and its bioderivatives, such as octyl acetate and octyl-β-D-glucoside (Sattayawat et al., 2020). *‘Uc*FatB1 was reported to be highly specific towards dodecanoic acid (C12:0) (Voelker et al., 1992) and has recently been combined with a SAM-dependent methyl transferase for bioproduction of methyl laurate in *Synechocystis* (Yunus et al., 2020). ‘*Ca*FatB3.5 and *‘ChoFatB2.2* only resulted in meager quantities of mainly 1-nonene (Fig. 4E). A more systematic approach was used in *‘Uc*FatB1 expressing strains, employing varying strength promoters and RBSs in front of the two plasmid-encoded genes. The solvent overlay of the best producing strain contained 231.07 mg/L (134.16 mg/g CDW) and 25.92 mg/L (15.05 mg/g CDW) of 1-undecene, and 1-tridecene, respectively (Fig. 4F).

### 3.4 Module 4 – Biosynthesis of C7-C13 alkanes with FAP

Previously, we demonstrated that a *Synechocystis* strain expressing truncated FAP and ‘tesA can efficiently convert C12-C18 free fatty acids into C11-C17 alka(e)nes (Yunus et al., 2018). However, all alkanes accumulated intracellularly. Here we sought to engineer *Synechocystis* for production of shorter alkanes in the hope that they would accumulate extracellularly. To achieve this, we combined either ‘*Uc*FatB1 or *Cp*FatB1.4 (Lozada et al., 2018; Sattayawat et al., 2020) with a fatty acid photodecarboxylase (FAP) from *Chlorella variabilis* (Fig. 5A). The cobalt-inducible Pcoa promoter was used to regulate the expression of *‘Cp*FatB1.4 whilst a constitutive Ptrc promoter was used to express FAP (Fig. 5B). Strains were grown under continuous illumination from warm white and blue LEDs required for cell growth and activation of the FAP enzyme, respectively (Fig. 5C). A range of C7-C13 alkanes accumulated extracellularly in the solvent overlay in strains expressing *Cp*FatB1.4 as the thioesterase (Fig. 5D). Liquid cultures induced with cobalt every two days produced higher alkane titer than those with a single dose of cobalt. In such latter cultures, the alkane titer decreased during the ten days of cultivation, suggesting that a portion, especially heptane, might have evaporated even in the presence of 10% (v/v) hexadecane solvent overlay. In strains expressing *‘Uc*FatB1 and FAP under a constitutive Ptrc promoter, varying the RBS sequence in front of the Fap-encoding gene enabled accumulation up to 15 mg/L undecane and 11 mg/L tridecane (Fig. 5E). Altogether, the experiments demonstrated that the combination of FAP with different thioesterases can enable the conversion of CO_2_ into alkanes that naturally excrete from the cell. Further improvements to the selectivity of FAP and environmental conditions are necessary to increase the partitioning of carbon into alkanes. Comparisons with UndB (Fig. 4F vs. Fig. 5E) suggest that FAP activity is limiting flux through the alkane pathway.

**Figure 5.**
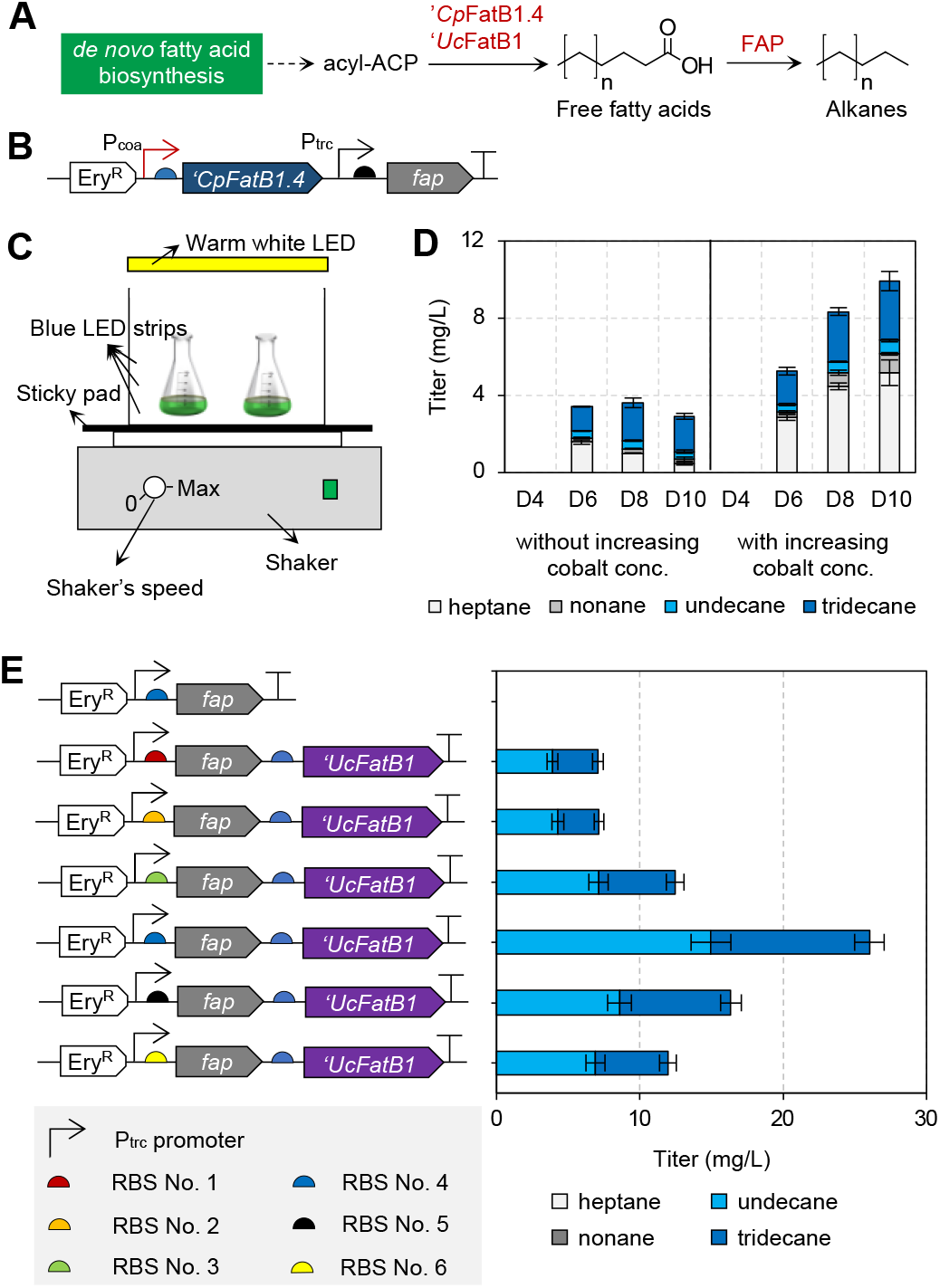
Production of short-medium chain length of alkanes. (A) Schematic diagram of alkane production. Two thioesterases (*‘Cp*FatB1.4 and *‘Uc*FatB1) were tested to convert acyl-ACPs into their corresponding free fatty acids. Free fatty acids were then decarboxylated to yield alkanes by a fatty acid photodecarboxylase (FAP). (B) Schematic diagram of plasmid that carries *‘Cp*FatB1.4 and FAP. (C) Experimental setup for alkane production in the presence of blue LED strips to activate the FAP enzyme. (D) Titer of alkanes from strains expressing *‘Cp*FatB1.4 and FAP (Fig. 5B) during ten days of cultivation with and without increasing cobalt inducer concentration. (E) Titer of alkanes from strains expressing FAP and *‘Uc*FatB1 with different RBSs. All liquid cultures were cultivated for two days with starting OD_730_ 0.2 and overlaid with 10% (v/v) hexadecane on day two. For strains shown in Fig. 5D, liquid cultures were induced with 625 nM cobalt on day two and repeated on day four and six for samples labelled as ‘increasing cobalt concentration’. Data were obtained from three biological replicates and error bars represent standard deviation.

## 4. Discussion

The direct sunlight-driven biological conversion of CO_2_ into engine-ready fuel is still only pursued in academic laboratories as far as we are aware. Whether or not it can be commercialized in the future remains to be seen and in the meanwhile it remains pertinent to pursue systems that are operationally optimal, ideally biocatalytic (i.e. >50% conversion of substrate). In fact, biocatalytic systems have already been achieved for some product (Angermayr et al., 2014; Liu et al., 2019), but more solutions are surely possible. In this study, we demonstrated that renewable production of excreted short to medium chain length hydrocarbons was possible via four different routes.

First, we established pathways for propane production by implementing a modified Clostridial pathway for *n*-butanol biosynthesis. To reduce the conversion of butyraldehyde to n-butanol, we introduced a CRISPRi system to repress the expression of three endogenous putative aldehyde reductases/dehydrogenases. This resulted in improved propane production. Most of the butyraldehyde substrate was still diverted into n-butanol, however. Additional unknown putative aldehyde reductases exist in *Synechocystis* that would be of interest to modify in future studies. The conversion of C4 precursor into propane in our best case was lower than 0.1%, whilst a similar pathway in *E. coli* achieved >50% acid-to-alkane conversion (Kallio et al., 2014). This is surprising given that Ado is native to cyanobacteria. However, Ado was also not particularly effective with longer (C14-C18) chain-length precursors in a previous study (Yunus et al., 2018). A major difference with the native alkane biosynthesis pathway in wild-type cyanobacteria is that the aldehyde precursor in our strains is provided by an introduced pathway. If Ado activity in cyanobacteria is dependent on specific localization and/or metabolon formation with Aar (Warui et al., 2015), then this might explain why Ado is reluctant to accept alternative sources of aldehyde.

For the second module, the aim was to produce pentane via a lipoxygenase pathway. Lipoxygenase are commonly found in many organisms, including cyanobacteria (Stolterfoht et al., 2019). Depending on the position of the C atom on fatty acids that become oxygenated, LOX enzymes can be classified into 5-, 8-, 9-, 10-, 11-, 12-, 13-, and 15-LOXs (Heshof et al., 2016). There are only a few cyanobacteria strains where LOX have been identified and characterized. For example, *Nostoc punctiforme* PCC 73102 and *Anabaena* sp. PCC7120 harbor 13-LOX (Koeduka et al., 2007; Lang and Feussner, 2007) and 9-LOX (Zhang et al., 2013, 2012), respectively, whilst *Cyanothece* sp. PCC 8801 harbors both 9-LOX (Newie et al., 2016) and 11-LOX (Andreou et al., 2010; Newie et al., 2017). A homology search with the Basic Local Alignment Search Tool (BLAST) (Johnson et al., 2008), using either plant (*Arabidopsis thaliana*; Q06327) or microbial LOX sequences (*Anabaena* sp. PCC 7120; WP_010994078, *Nostoc punctiforme* PCC 73102; ZP_00106490 and ZP_00107030) as input and a 0.05 threshold cutoff resulted in no hits. Additionally, to find if there are native lipoxygenases that form interaction/association with desaturases genes, we also searched a text mining network based on species-dependent protein-protein or transcriptome interactions (Kreula et al., 2018). The expanded network of desC or desA did not show protein or transcriptome association with native putative lipoxygenase (Supplementary Figure S5). Altogether, this may indicate that *Synechocystis* sp. PCC 6803 lacks lipoxygenase enzymes, or that they still remain to be discovered.

For the third and fourth modules, we combined C8-C12 thioesterases with previously studied decarboxylases that natively produce hydrocarbons directly from free fatty acids. Fortunately, the majority of the biosynthesised C11 hydrocarbons were found to accumulate in the solvent overlay fraction suggesting it can naturally be excreted as long as there is a hydrophobic space nearby. The difference between strains over-expressing FAP (Fig. 5E) vs. UndB (Fig. 4F) was striking even though both utilised ‘*Uc*FatB1. Both the total accumulation and the chain-length profile varied, suggesting that UndB functions far better with C12 fatty acids and that the native FAP used in this study displays limited activity towards C12. Noticeable variation in productivity was also observed in response to changes in the construct design. Although the quantity of key proteins was not measured, it highlights the importance of molecular construct optimization.

## 5. Conclusion

The main objective with this study was to investigate options for the direct conversion of CO_2_ into engine-ready fuels that are naturally secreted. Synthetic metabolic pathways enabling the biosynthesis of several such molecules were implemented and found to result in external accumulation when cultured in the presence of a solvent overlay, or in closed serum bottles. This opens up the opportunity to milk photosynthetic microorganisms for products in continuous cultivation systems. Although not exhaustive, the reported experiments indicate that UndB remains the best terminal enzyme for hydrocarbon biosynthesis in *Synechocystis* for the C12 precursor chain-length, whilst this and previous studies suggest that FAP is superior to Ado for alkane biosynthesis with all chain-lengths tested so far.

## Acknowledgement

This project has received funding from the European Union’s Horizon 2020 research and innovation program project PHOTOFUEL under the grant agreement No. <u>640720</u> (PRJ), a PhD scholarship from the Indonesia Endowment Fund for Education (LPDP)(ISY) and the Swedish Research Council (2016-06160)(EPH).

## References

Albers, S.C., Gallegos, V.A., Peebles, C.A., 2015. Engineering of genetic control tools in Synechocystis sp. PCC 6803 using rational design techniques. J Biotechnol 216. https://doi.org/10.1016/j.jbiotec.2015.09.042

Amer, M., Wojcik, E.Z., Sun, C., Hoeven, R., Hughes, J.M.X., Faulkner, M., Yunus, I.S., Tait, S., Johannissen, L.O., Hardman, S.J.O., Heyes, D.J., Chen, G.-Q., Smith, M.H., Jones, P.R., Toogood, H.S., Scrutton, N.S., 2020. Low carbon strategies for sustainable bio-alkane gas production and renewable energy. Energy Environ. Sci. https://doi.org/10.1039/D0EE00095G

Andreou, A., Göbel, C., Hamberg, M., Feussner, I., 2010. A Bisallylic Mini-Lipoxygenase from Cyanobacterium Cyanothece Sp. That Has an Iron as Cofactor. J. Biol. Chem. 285, 14178.

Anfelt, J., Kaczmarzyk, D., Shabestary, K., Renberg, B., Rockberg, J., Nielsen, J., Uhlén, M., Hudson, E.P., 2015. Genetic and nutrient modulation of acetyl-CoA levels in Synechocystis for n-butanol production. Microb. Cell Fact. 14, 167. https://doi.org/10.1186/s12934-015-0355-9

Angermayr, S.A., Van Der Woude, A.D., Correddu, D., Vreugdenhil, A., Verrone, V., Hellingwerf, K.J., 2014. Exploring metabolic engineering design principles for the photosynthetic production of lactic acid by Synechocystis sp. PCC6803. Biotechnol. Biofuels 7, 1–15. https://doi.org/10.1186/1754-6834-7-99

Atsumi, S., Cann, A.F., Connor, M.R., Shen, C.R., Smith, K.M., Brynildsen, M.P., Chou, K.J.Y., Hanai, T., Liao, J.C., 2008. Metabolic engineering of Escherichia coli for 1-butanol production. Metab. Eng. 10, 305–311. https://doi.org/10.1016/j.ymben.2007.08.003

Bikard, D., Jiang, W., Samai, P., Hochschild, A., Zhang, F., Marraffini, L. a., 2013. Programmable repression and activation of bacterial gene expression using an engineered CRISPR-Cas system. Nucleic Acids Res. 41, 7429.

Blazeck, J., Liu, L., Knight, R., Alper, H.S., 2013. Heterologous production of pentane in the oleaginous yeast Yarrowia lipolytica. J. Biotechnol. 165, 184–194. https://doi.org/10.1016/j.jbiotec.2013.04.003

Chen, G., Qu, S., Wang, Q., Bian, F., Peng, Z., Zhang, Y., Ge, H., Yu, J., Xuan, N., Bi, Y., He, Q., 2014. Transgenic expression of delta-6 and delta-15 fatty acid desaturases enhances omega-3 polyunsaturated fatty acid accumulation in Synechocystissp. PCC6803. Biotechnol. Biofuels 7, 32. https://doi.org/10.1186/1754-6834-7-32

Donovan, D.H., Menzel, D.B., 1978. Mechanisms of lipid peroxidation: Iron catalyzed decomposition of fatty acid hydroperoxides as the basis of hydrocarbon evolution in vivo. Experientia 34, 775–776. https://doi.org/10.1007/BF01947320

Fasaei, F., Bitter, J.H., Slegers, P.M., van Boxtel, A.J.B., 2018. Techno-economic evaluation of microalgae harvesting and dewatering systems. Algal Res. 31, 347–362. https://doi.org/10.1016/j.algal.2017.11.038

Feussner, I., Wasternack, C., 2002. The lipoxygenase pathway. Annu. Rev. Plant Biol. 53, 275–297. https://doi.org/10.1146/annurev.arplant.53.100301.135248

Gao, Z., Zhao, H., Li, Z., Tan, X., Lu, X., 2012. Photosynthetic production of ethanol from carbon dioxide in genetically engineered cyanobacteria. Energy Env. Sci 5. https://doi.org/10.1039/C2EE22675H

Garssen, G.J., Vliegenthart, J.F.G., Boldingh, J., 1971. An anaerobic reaction between lipoxygenase, linoleic acid and its hydroperoxides. Biochem. J. 122, 327–332. https://doi.org/10.1042/bj1220327

Heshof, R., de Graaff, L.H., Villaverde, J.J., Silvestre, A.J.D., Haarmann, T., Dalsgaard, T.K., Buchert, J., 2016. Industrial potential of lipoxygenases. Crit. Rev. Biotechnol. 36, 665–674. https://doi.org/10.3109/07388551.2015.1004520

Hintzpeter, J., Martin, H.-J., Maser, E., 2015. Reduction of lipid peroxidation products and advanced glycation end-product precursors by cyanobacterial aldo-keto reductase AKR3G1--a founding member of the AKR3G subfamily. FASEB J. 29, 263.

Horvat, R.J., Lane, W.G., Ng, H., Shepherd, A.D., 1964. Saturated Hydrocarbons from Autoxidizing Methyl Linoleate. Nature 203, 523–524. https://doi.org/10.1038/203523b0

Huang, H.H., Lindblad, P., 2013. Wide-dynamic-range promoters engineered for cyanobacteria. J Biol Eng 7. https://doi.org/10.1186/1754-1611-7-10

Johnson, M., Zaretskaya, I., Raytselis, Y., Merezhuk, Y., McGinnis, S., Madden, T.L., 2008. NCBI BLAST: a better web interface. Nucleic Acids Res. 36, W5–W9. https://doi.org/10.1093/nar/gkn201

Kaczmarzyk, D., Fulda, M., 2010. Fatty Acid Activation in Cyanobacteria Mediated by Acyl-Acyl Carrier Protein Synthetase Enables Fatty Acid Recycling. Plant Physiol. 152, 1598–1610. https://doi.org/10.1104/pp.109.148007

Kallio, P., Pásztor, A., Thiel, K., Akhtar, M.K., Jones, P.R., 2014. An engineered pathway for the biosynthesis of renewable propane. Nat. Commun. 5, 4731.

Khara, B., Menon, N., Levy, C., Mansell, D., Das, D., Marsh, E.N.G., Leys, D., Scrutton, N.S., 2013. Production of Propane and Other Short-Chain Alkanes by Structure-Based Engineering of Ligand Specificity in Aldehyde-Deformylating Oxygenase. ChemBioChem 14, 1204–1208. https://doi.org/10.1002/cbic.201300307

Kim, J., Yoo, G., Lee, H., Lim, J., Kim, K., Kim, C.W., Park, M.S., Yang, J.-W., 2013. Methods of downstream processing for the production of biodiesel from microalgae. Biotechnol. Adv. 31, 862–876. https://doi.org/10.1016/j.biotechadv.2013.04.006

Koeduka, T., Kajiwara, T., Matsui, K., 2007. Cloning of Lipoxygenase Genes from a Cyanobacterium, Nostoc punctiforme, and Its Expression in Eschelichia coli. Curr. Microbiol. 54, 315.

Kreula, S.M., Kaewphan, S., Ginter, F., Jones, P.R., 2018. Finding novel relationships with integrated gene-gene association network analysis of Synechocystis sp. PCC 6803 using species-independent text-mining. PeerJ 6, e4806. https://doi.org/10.7717/peerj.4806

Krivoruchko, A., Serrano-Amatriain, C., Chen, Y., Siewers, V., Nielsen, J., 2013. Improving biobutanol production in engineered Saccharomyces cerevisiae by manipulation of acetyl-CoA metabolism. J. Ind. Microbiol. Biotechnol. 40, 1051–1056. https://doi.org/10.1007/s10295-013-1296-0

Lan, E.I., Liao, J.C., 2012. ATP drives direct photosynthetic production of 1-butanol in cyanobacteria. Proc. Natl. Acad. Sci. 109, 6018–6023. https://doi.org/10.1073/pnas.1200074109

Lan, E.I., Liao, J.C., 2011. Metabolic engineering of cyanobacteria for 1-butanol production from carbon dioxide. Metab Eng 13. https://doi.org/10.1016/j.ymben.2011.04.004

Lan, E.I., Ro, S.Y., Liao, J.C., 2013. Oxygen-tolerant coenzyme A-acylating aldehyde dehydrogenase facilitates efficient photosynthetic n-butanol biosynthesis in cyanobacteria. Energy Environ. Sci. 6, 2672. https://doi.org/10.1039/c3ee41405a

Lang, I., Feussner, I., 2007. Oxylipin Formation in Nostoc punctiforme (PCC73102). Phytochemistry 68, 1120.

Larson, M.H., Gilbert, L.A., Wang, X., Lim, W.A., Weissman, J.S., Qi, L.S., 2013. CRISPR interference (CRISPRi) for sequence-specific control of gene expression. Nat. Protoc. 8, 2180.

Li, J., Ma, Y., Liu, N., Eser, B.E., Guo, Z., Jensen, P.R., Stephanopoulos, G., 2020. Synthesis of high-titer alka(e)nes in Yarrowia lipolytica is enabled by a discovered mechanism. Nat. Commun. 11, 6198. https://doi.org/10.1038/s41467-020-19995-0

Liu, X., Miao, R., Lindberg, P., Lindblad, P., 2019. Modular engineering for efficient photosynthetic biosynthesis of 1-butanol from CO2 in cyanobacteria. Energy Environ. Sci. 12, 2765–2777. https://doi.org/10.1039/C9EE01214A

Lozada, N.J.H., Lai, R.-Y., Simmons, T.R., Thomas, K.A., Chowdhury, R., Maranas, C.D., Pfleger, B.F., 2018. Highly Active C8-Acyl-ACP Thioesterase Variant Isolated by a Synthetic Selection Strategy. ACS Synth. Biol. 7, 2205–2215. https://doi.org/10.1021/acssynbio.8b00215

Menon, N., Pásztor, A., Menon, B.R.K., Kallio, P., Fisher, K., Akhtar, M.K., Leys, D., Jones, P.R., Scrutton, N.S., 2015. A microbial platform for renewable propane synthesis based on a fermentative butanol pathway. Biotechnol. Biofuels 8, 61. https://doi.org/10.1186/s13068-015-0231-1

Miao, R., Liu, X., Englund, E., Lindberg, P., Lindblad, P., 2017. Isobutanol production in Synechocystis PCC 6803 using heterologous and endogenous alcohol dehydrogenases. Metab. Eng. Commun. 5, 45–53. https://doi.org/10.1016/j.meteno.2017.07.003

Newie, J., Andreou, A., Neumann, P., Einsle, O., Feussner, I., Ficner, R., 2016. Crystal Structure of a Lipoxygenase from Cyanothece Sp. May Reveal Novel Features for Substrate Acquisition. J. Lipid Res. 57, 276.

Newie, J., Neumann, P., Werner, M., Mata, R.A., Ficner, R., Feussner, I., 2017. Lipoxygenase 2 from Cyanothece Sp. Controls Dioxygen Insertion by Steric Shielding and Substrate Fixation. Sci. Rep. 7, 2069.

Nielsen, D.R., Leonard, E., Yoon, S.-H., Tseng, H.-C., Yuan, C., Prather, K.L.J., 2009. Engineering alternative butanol production platforms in heterologous bacteria. Metab. Eng. 11, 262–273. https://doi.org/10.1016/j.ymben.2009.05.003

Ögmundarson, Ó., Herrgård, M.J., Forster, J., Hauschild, M.Z., Fantke, P., 2020. Addressing environmental sustainability of biochemicals. Nat. Sustain. 3, 167–174. https://doi.org/10.1038/s41893-019-0442-8

Perin, G., Jones, P.R., 2019. Economic feasibility and long-term sustainability criteria on the path to enable a transition from fossil fuels to biofuels. Curr. Opin. Biotechnol. 57, 175–182. https://doi.org/10.1016/j.copbio.2019.04.004

Qi, L.S., Larson, M.H., Gilbert, L.A., Doudna, J.A., Weissman, J.S., Arkin, A.P., Lim, W.A., 2013. Repurposing CRISPR as an RNA-guided platform for sequence-specific control of gene expression. Cell 152, 1173.

Rui, Z., Harris, N.C., Zhu, X., Huang, W., Zhang, W., 2015. Discovery of a Family of Desaturase-Like Enzymes for 1-Alkene Biosynthesis. ACS Catal. 5, 7091–7094. https://doi.org/10.1021/acscatal.5b01842

Sanders, T.H., Pattee, H.E., Singleton, J.A., 1975. Aerobic pentane production by soybean lipoxygenase isozymes. Lipids 10, 568–570. https://doi.org/10.1007/BF02532364

Santos-Merino, M., Garcillán-Barcia, M.P., de la Cruz, F., 2018. Engineering the fatty acid synthesis pathway in Synechococcus elongatus PCC 7942 improves omega-3 fatty acid production. Biotechnol. Biofuels 11, 239. https://doi.org/10.1186/s13068-018-1243-4

Sattayawat, P., Yunus, I.S., Jones, P.R., 2020. Bioderivatization as a concept for renewable production of chemicals that are toxic or poorly soluble in the liquid phase. Proc. Natl. Acad. Sci. 117, 1404 LP–1413. https://doi.org/10.1073/pnas.1914069117

Schirmer, A., Rude, M.A., Li, X., Popova, E., del Cardayre, S.B., 2010. Microbial biosynthesis of alkanes. Science (80-.). 329, 559.

Sheppard, M.J., Kunjapur, A.M., Prather, K.L.J., 2016. Modular and selective biosynthesis of gasoline-range alkanes. Metab. Eng. 33, 28–40. https://doi.org/10.1016/j.ymben.2015.10.010

Sorigué, D., Légeret, B., Cuiné, S., Blangy, S., Moulin, S., Billon, E., Richaud, P., Brugière, S., Couté, Y., Nurizzo, D., Müller, P., Brettel, K., Pignol, D., Arnoux, P., Li-Beisson, Y., Peltier, G., Beisson, F., 2017. An algal photoenzyme converts fatty acids to hydrocarbons. Science (80-.). 357, 903 LP–907. https://doi.org/10.1126/science.aan6349

Steen, E.J., Chan, R., Prasad, N., Myers, S., Petzold, C.J., Redding, A., Ouellet, M., Keasling, J.D., 2008. Metabolic engineering of Saccharomyces cerevisiae for the production of n-butanol. Microb. Cell Fact. 7, 36. https://doi.org/10.1186/1475-2859-7-36

Stolterfoht, H., Rinnofner, C., Winkler, M., Pichler, H., 2019. Recombinant Lipoxygenases and Hydroperoxide Lyases for the Synthesis of Green Leaf Volatiles. J. Agric. Food Chem. 67, 13367–13392. https://doi.org/10.1021/acs.jafc.9b02690

Storch, M., Casini, A., Mackrow, B., Fleming, T., Trewhitt, H., Ellis, T., Baldwin, G.S., 2015. BASIC: A New Biopart Assembly Standard for Idempotent Cloning Provides Accurate, Single-Tier DNA Assembly for Synthetic Biology. ACS Synth. Biol. 4, 781–787. https://doi.org/10.1021/sb500356d

Trautmann, D., Beyer, P., Al-Babili, S., 2013. The ORF slr0091 of Synechocystis sp. PCC6803 encodes a high-light induced aldehyde dehydrogenase converting apocarotenals and alkanals. FEBS J. 280, 3685.

Voelker, T.A., Worrell, A.C., Anderson, L., Bleibaum, J., Fan, C., Hawkins, D.J., Radke, S.E., Davies, H.M., 1992. Fatty acid biosynthesis redirected to medium chains in transgenic oilseed plants. Science (80-.). 257, 72 LP–74. https://doi.org/10.1126/science.1621095

Warui, D.M., Pandelia, M.-E., Rajakovich, L.J., Krebs, C., Bollinger, J.M., Booker, S.J., 2015. Efficient Delivery of Long-Chain Fatty Aldehydes from the Nostoc punctiforme Acyl-Acyl Carrier Protein Reductase to Its Cognate Aldehyde-Deformylating Oxygenase. Biochemistry 54, 1006–1015. https://doi.org/10.1021/bi500847u

Yao, L., Cengic, I., Anfelt, J., Hudson, E.P., 2016. Multiple Gene Repression in Cyanobacteria Using CRISPRi. ACS Synth. Biol. 5, 207–212. https://doi.org/10.1021/acssynbio.5b00264

Yoshino, T., Kakunaka, N., Liang, Y., Ito, Y., Maeda, Y., Nomaguchi, T., Matsunaga, T., Tanaka, T., 2017. Production of ω3 fatty acids in marine cyanobacterium Synechococcus sp. strain NKBG 15041c via genetic engineering. Appl. Microbiol. Biotechnol. 101, 6899–6905. https://doi.org/10.1007/s00253-017-8407-1

Yunus, I.S., Jones, P.R., 2018. Photosynthesis-dependent biosynthesis of medium chain-length fatty acids and alcohols. Metab. Eng. 49. https://doi.org/10.1016/j.ymben.2018.07.015

Yunus, I.S., Palma, A., Trudeau, D.L., Tawfik, D.S., Jones, P.R., 2020. Methanol-free biosynthesis of fatty acid methyl ester (FAME) in Synechocystis sp. PCC 6803. Metab. Eng. 57, 217–227. https://doi.org/10.1016/j.ymben.2019.12.001

Yunus, I.S., Wang, Z., Sattayawat, P., Muller, J., Zemichael, F.W., Hellgardt, K., Jones, P.R., 2021. Improved Bioproduction of 1-Octanol Using Engineered Synechocystis sp. PCC 6803. ACS Synth. Biol. 10, 1417–1428. https://doi.org/10.1021/acssynbio.1c00029

Yunus, I.S., Wichmann, J., Wördenweber, R., Lauersen, K.J., Kruse, O., Jones, P.R., 2018. Synthetic metabolic pathways for photobiological conversion of CO2 into hydrocarbon fuel. Metab. Eng. 49, 201–211. https://doi.org/10.1016/j.ymben.2018.08.008

Zhang, C., Tao, T., Ying, Q., Zhang, D., Lu, F., Bie, X., Lu, Z., 2012. Extracellular Production of Lipoxygenase from Anabaena Sp. PCC 7120 in Bacillus subtilis and Its Effect on Wheat Protein. Appl. Microbiol. Biotechnol. 94, 949.

Zhang, C., Zhang, S., Lu, Z., Bie, X., Zhao, H., Wang, X., Lu, F., 2013. Effects of Recombinant Lipoxygenase on Wheat Flour, Dough and Bread Properties. Food Res. Int. 54, 26.

Zhu, Z., Zhou, Y.J., Kang, M.-K., Krivoruchko, A., Buijs, N.A., Nielsen, J., 2017. Enabling the synthesis of medium chain alkanes and 1-alkenes in yeast. Metab. Eng. 44, 81–88. https://doi.org/10.1016/j.ymben.2017.09.007

