## Supplementary material for "Synthetic metabolic pathways for conversion of CO_2_ into secreted short-to medium-chain hydrocarbons using cyanobacteria"

### Supplementary Note S1. Design of sgRNA

Design of 1-sgRNA

tccctatcagtgatagagattgacatccctatcagtgatagatataatgtgtggtaccggggtcgtattcaaagggtgttttagagctagaaatag
caagttaaaataaggctagtccgttatcaacttgaaaaagtggcaccgagtcggtgcttttttta

Design of 4-sgRNAs

tccctatcagtgatagagattgacatccctatcagtgatagatataatgtgtggtatctgctccccattgctcaaagttttagagctagaaatagca
agttaaaataaggctagtccgttatcaacttgaaaaagtggcaccgagtcggtgctttttttttactagagtccctatcagtgatagagattgacat
ccctatcagtgatagatataatgtgtggtacttcaaaggcaacggcggcacgttttagagctagaaatagcaagttaaaataaggctagtccgtt
atcaacttgaaaaagtggcaccgagtcggtgctttttttttactagagtccctatcagtgatagagattgacatccctatcagtgatagatataatgt
gtggtatttcgtttacatagctttcaaaccgttttagagctagaaatagcaagttaaaataaggctagtccgttatcaacttgaaaaagtggcacc
gagtcggtgctttttttttactagagtccctatcagtgatagagattgacatccctatcagtgatagatataatgtgtggtaccggggtcgtattcaa
agggtgttttagagctagaaatagcaagttaaaataaggctagtccgttatcaacttgaaaaagtggcaccgagtcggtgctttttttttactagag

Legend:

L31 promoter

slr0942 target

sll0990 target

slr0091 target

slr1192 target

dCas9-binding

Rho-independent terminator

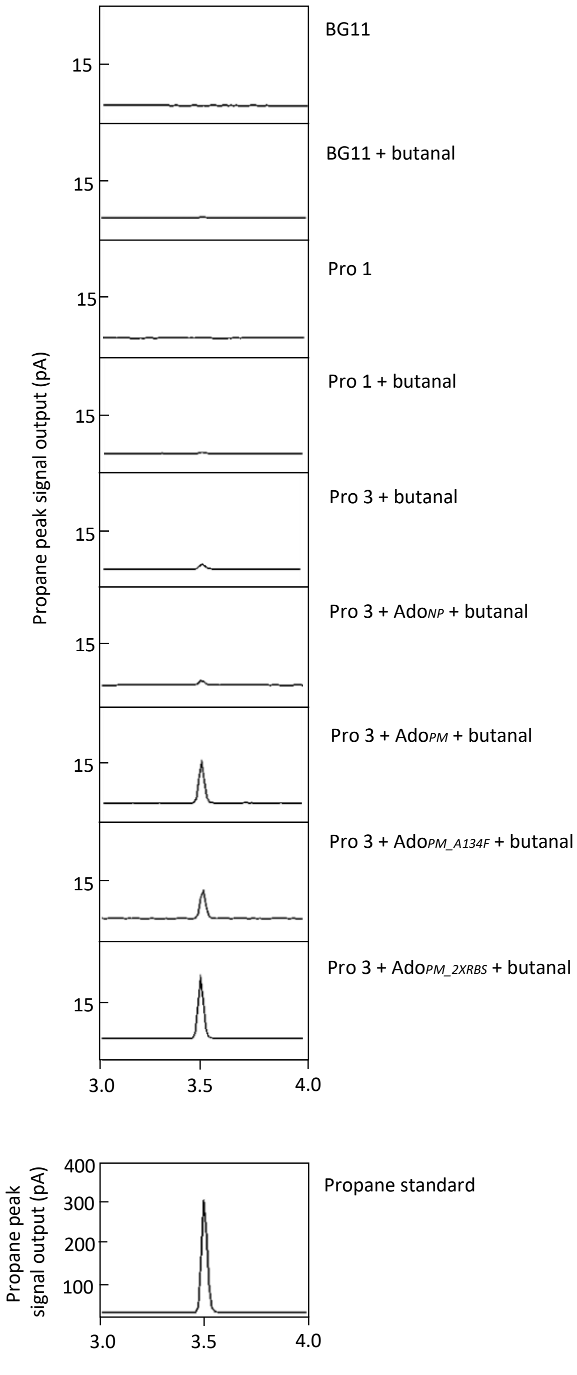

**Supplementary Figure S1. Selection of superior aldehyde deformylating oxygenase by butanal feeding**

Propane was sampled from the headspace and analyzed using GC-FID.

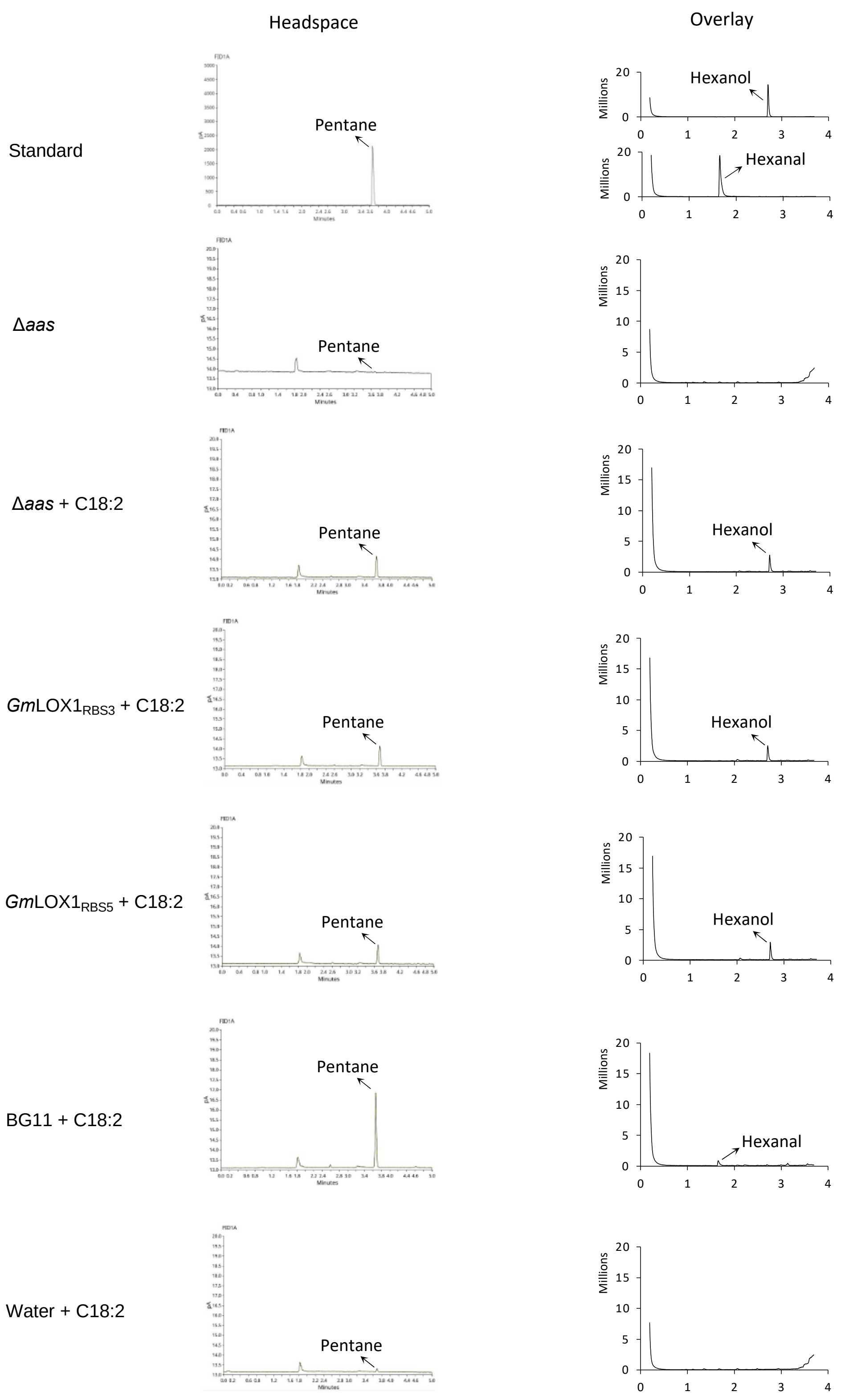

**Supplementary Figure S2. Chromatograms of pentane, 1-hexanol, and hexanal obtained from samples shown in Fig. 3D-F**

Pentane was sampled from the headspace and analyzed using GC-FID. Hexanal and 1-hexanol were sampled from the solvent overlay and analyzed using GC-MS.

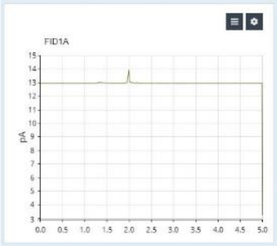

Δ*aas*

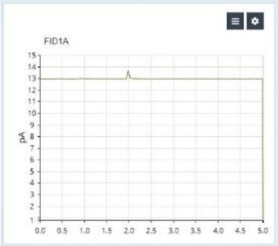

Δ*aas* + butyric acid (2 µL)

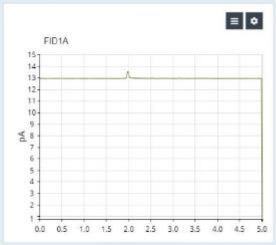

Δ*aas* + Ptrc-UndB

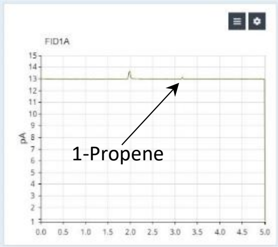

Δ*aas* + Ptrc-UndB + butyric acid (2 µL)

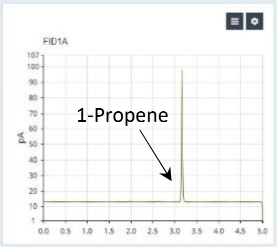

1-Propene standard (174 ug/L)

**Supplementary Figure S3. Biosynthesis of 1-Propene (propylene)**

Production of 1-Propene was achieved by expressing UndB in a *Synechocystis* sp. PCC 6803 strain lacking acyl-ACP synthetase (*aas*) in the presence of exogenously added butyric acid. 1-Propene was analyzed using GC-FID as follows: 500 µL of headspace sample was taken and analyzed using a 8860 Gas Chromatograph (GC) System equipped with a Flame Ionization Detector (FID) and a GS-GasPro UST 1446134 column (Agilent, 113-4332). The inlet temperature was set at 250 $^{o}$C (8.7 psi). Flow rate was maintained at 22.143 ml/min, with 0.743 ml/min purge flow. Oven was set at 120 $^{o}$C and hold for 5 min.

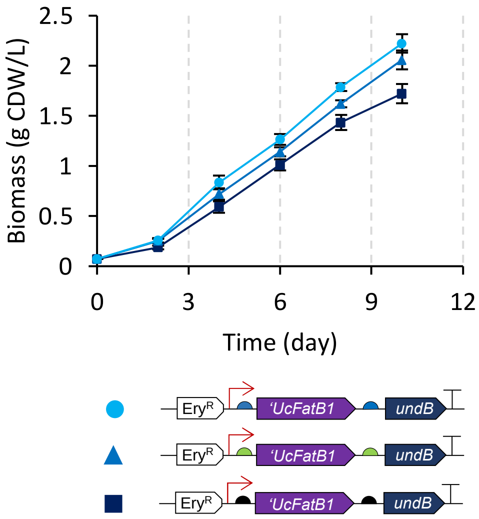

**Supplementary Figure S4. Growth profile of some of strains shown in Fig. 4F**

$OD_{730}$ was measured every two days using a spectrophotometer (Tecan Infinite 200 PRO). By using conversion $OD_{730}$ 1 = 0.36 g CDW/L, the value of $OD_{730}$ was converted into biomass (g CDW/L) and used to calculate carbon partitioning to 1-undecene and 1-tridecene.

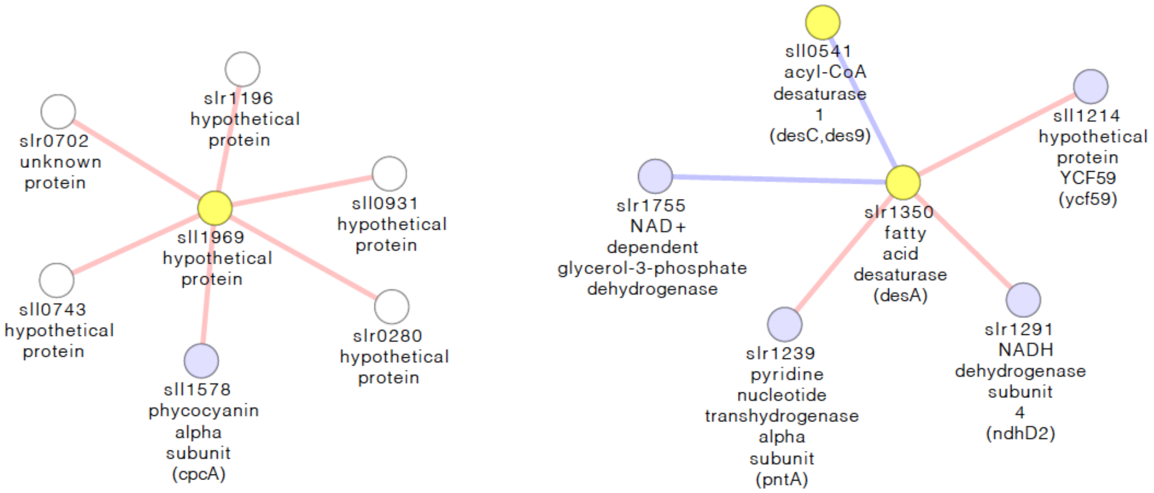

**Supplementary Figure S5. Proteome or transcriptome association of expanded network of lipA (sll1969) and desA (slr1350)**. The initial three-node selection (yellow) was expanded through existing edges with the "First neighbours of selected nodes (undirected)'' function of Cytoscape.

**Supplementary Table S1. The amino acid sequences of all enzymes used in this study**

| Enzyme | Amino acid sequence |
|---|---|
| NphT7 | MHHHHHHTDVRFRIIGTGAYVPERIVSNDEVGAPAGVDDDWITRKTGIRQRR<br>WAADDQATSDLATAAGRAALKAAGITPEQLTVIAVATSTPDRPQPPTAAYVQH<br>HLGATGTAAFDVNAVCSGTVFALSSVAGTLVYRGGYALVIGADLYSRILNPADRK<br>TVVLFGDGAGAMVLGPTSTGTGPIVRRVALHTFGGLTDLIRVPAGGSRQPLDTD<br>GLDAGLQYFAMDGREVRRFVTEHLPQLIKGFLHEAGVDAADISHFVPHQANGV<br>MLDEVFGELHLPRATMHRTVETYGNTGAASIPITMDAAVRAGSFRPGELVLLAG<br>FGGGMAASFALIEW* |
| XfpK | MHHHHHHTNPVIGTPWQKLDRPVSEEAIEGMDKYWRVTNYMSIGQIYLRSNP<br>LMKEPFTRDDVKHRLVGHWGTTPGLNFLLAHINRLIADHQQNTVFIMGPGHG<br>GPAGTSQSYVDGTYTEYYPNITKDEAGLQKFFRQFSYPGGIPSHFAPETPGSIHEG<br>GELGYALSHAYGAVMNNPSLFVPCIIGDGEAETGPLATGWQSNKLVNPRTDGIV<br>LPILHLNGYKIANPTILARISDEELHDFFRGMGYHPYEFVAGFDNEDHMSIHRRFA<br>ELFETIFDEICDIKAAAQTDDMTRPFYPMLIFRTPKGWTCPKFIDGKKTEGSWRA<br>HQVPLASARDTEEHFEVLKGWMESYKPEELFNADGSIKDDVTAFMPKGELRIGA<br>NPNANGGVIREDLKLPELDQYEVTGVKEYGHGWGQVEAPRALGAYCRDIIKNN<br>PDSFRIFGPDETASNRLNATYEVTDKQWDNGYLSGLVDEHMAVTGQVTEQLSE<br>HQCEGFLEAYLLTGRHGIWSSYESFVHVIDSMLNQHAKWLEATVREIPWRKPISS<br>VNLLVSSHVWRQDHNGFSHQDPGVTSLLINKTFNNDHVTNIYFATDANMLLAIS<br>EKCFKSTNKINAIFAGKQPAPTWVTLDEARAELEAGAAEWKWASNAENNDEV<br>QVVLASAGDVPTQELMAASDALNKMGIKFKVVNVVDLLKLQSRENNDEALTDE<br>EFTELFTADKPVLFAYHSYAQDVRGLIYDRPNHDNFHVVGYKEQGSTTTPFDMV<br>RVNDMDRYALQAAALKLIDADKYADKIDELNAFRKKAFQFAVDNGYDIPEFTD<br>WVYPDVKVDETQMLSATAATAGDNE* |
| PduP | MEQKLISEEDLNTSELETLIRTILSEQLTTPAQTPVQPQGKGIFQSVSEAIDAAHQA<br>FLRYQQCPLKTRSAIISAMRQELTPLLATLAEESANETGMGNKEDKFLKNKAALD<br>NTPGVEDLTTTALTGDGGMVLFEYSPFGVIGSVAPSTNPTETIINNSISMLAAGN<br>SVYFSPHPGAKKVSLKLISLIEEIAFRCCGIRNLVVTVAEPTFEATQQMMAHPRIA<br>VLAITGGPGIVAMGMKSGKKVIGAGAGNPPCIVDETADLVKAAEDIINGASFDY<br>NLPCIAEKSLIVVESVAERLVQQMQTFGALLLSPTDTDKLRAVCLPEGQANKKLV<br>GKSPSAMLEAAGIAVPAKAPRLLIAVVNADDPWVTSEQLMPMLPVVKVSDFDS<br>ALALALKVEEGLHHTAIMHSQNVSRLNLAARTLQTSIFVKNGPSYAGIGVGGEGF<br>TTFTIATPTGEGTTSARTFARSRRCVLTNGFSIR* |

| | |
|---|---|
| PhaJ | MEQKLISEEDLGGSSSAQSLEVGQKARLSKRFGAAEVAAFAALSEDFNPLHLDPA<br>FAATTAFERPIVHGMLLASLFSGLLGQQLPGKGSIYLGQSLSFKLPVFVGDEVTAE<br>VEVTALREDKPIATLTTRIFTQGGALAVTGEAVVKLP* |
| Ter | MDYKDDDDKGGSSIVKPMVRNNICLNAHPQGCKKGVEDQIEYTKKRITAEVKA<br>GAKAPKNVLVLGCSNGYGLASRITAAFGYGAATIGVSFEKAGSETKYGTPGWYN<br>NLAFDEAAKREGLYSVTIDGDAFSDEIKAQVIEEAKKKGIKFDLIVYSLASPVRTDP<br>DTGIMHKSVLKPFGKTFTGKTVDPFTGELKEISAEPANDEEAAATVKVMGGED<br>WERWIKQLSKEGLLEEGCITLAYSYIGPEATQALYRKGTIGKAKEHLEATAHRLNK<br>ENPSIRAFVSVNKGLVTRASAVIPVIPLYLASLFKVMKEKGNHEGCIEQITRLYAER<br>LYRKDGTIPVDEENRIRIDDWELEEDVQKAVSALMEKVTGENAESLTDLAGYRH<br>DFLASNGFDVEGINYEAEVERFDRI* |
| dCas9 | MDKKYSIGLAIGTNSVGWAVITDEYKVPSKKFKVLGNTDRHSIKKNLIGALLFDSG<br>ETAEATRLKRTARRRYTRRKNRICYLQEIFSNEMAKVDDSFFHRLEESFLVEEDKK<br>HERHPIFGNIVDEVAYHEKYPTIYHLRKKLVDSTDKADLRLIYLALAHMIKFRGHFL<br>IEGDLNPDNSDVDKLFIQLVQTYNQLFEENPINASGVDAKAILSARLSKSRRLENLI<br>AQLPGEKKNGLFGNLIALSLGLTPNFKSNFDLAEDAKLQLSKDTYDDDLDNLLAQI<br>GDQYADLFLAAKNLSDAILLSDILRVNTEITKAPLSASMIKRYDEHHQDLTLLKALV<br>RQQLPEKYKEIFFDQSKNGYAGYIDGGASQEEFYKFIKPILEKMDGTEELLVKLNR<br>EDLLRKQRTFDNGSIPHQIHLGELHAILRRQEDFYPFLKDNREKIEKILTFRIPYYVG<br>PLARGNSRFAWMTRKSEETITPWNFEEVVDKGASAQSFIERMTNFDKNLPNEK<br>VLPKHSLLYEYFTVYNELTKVKYVTEGMRKPAFLSGEQKKAIVDLLFKTNRKVTVK<br>QLKEDYFKKIECFDSVEISGVEDRFNASLGTYHDLLKIIKDKDFLDNEENEDILEDIV<br>LTLTLFEDREMIEERLKTYAHLFDDKVMKQLKRRRYTGWGRLSRKLINGIRDKQS<br>GKTILDFLKSDGFANRNFMQLIHDDSLTFKEDIQKAQVSGQGDSLHEHIANLAGS<br>PAIKKGILQTVKVVDELVKVMGRHKPENIVIEMARENQTTQKGQKNSRERMKRI<br>EEGIKELGSQILKEHPVENTQLQNEKLYLYYLQNGRDMYVDQELDINRLSDYDVD<br>AIVPQSFLKDDSIDNKVLTRSDKNRGKSDNVPSEEVVKKMKNYWRQLLNAKLIT<br>QRKFDNLTKAERGGLSELDKAGFIKRQLVETRQITKHVAQILDSRMNTKYDENDK<br>LIREVKVITLKSKLVSDFRKDFQFYKVREINNYHHAHDAYLNAVVGTALIKKYPKLE<br>SEFVYGDYKVYDVRKMIAKSEQEIGKATAKYFFYSNIMNFFKTEITLANGEIRKRPL<br>IETNGETGEIVWDKGRDFATVRKVLSMPQVNIVKKTEVQTGGFSKESILPKRNSD<br>KLIARKKDWDPKKYGGFDSPTVAYSVLVVAKVEKGKSKKLKSVKELLGITIMERSS<br>FEKNPIDFLEAKGYKEVKKDLIIKLPKYSLFELENGRKRMLASAGELQKGNELALPS<br>KYVNFLYLASHYEKLKGSPEDNEQKQLFVEQHKHYLDEIIEQISEFSKRVILADANL<br>DKVLSAYNKHRDKPIREQAENIIHLFTLTNLGAPAAFKYFDTTIDRKRYTSTKEVLD<br>ATLIHQSITGLYETRIDLSQLGGDEQKLISEEDL* |

| | |
|---|---|
| Ado$_{Pm}$ | MPTLEMPVAAVLDSTVGSSEALPDFTSDRYKDAYSRINAIVIEGEQEAHDNYIAIG<br>TLLPDHVEELKRLAKMEMRHKKGFTACGKNLGVEADMDFAREFFAPLRDNFQT<br>ALGQGKTPTCLLIQALLIEAFAISAYHTYIPVSDPFARKITEGVVKDEYTHLNYGEA<br>WLKANLESCREELLEANRENLPLIRRMLDQVAGDAAVLQMDKEDLIEDFLIAYQ<br>ESLTEIGFNTREITRMAAAALVS* |
| Ado$_{Np}$ | MAHHHHHHQQLTDQSKELDFKSETYKDAYSRINAIVIEGEQEAHENYITLAQLLP<br>ESHDELIRLSKMESRHKKGFEACGRNLAVTPDLQFAKEFFSGLHQNFQTAAAEG<br>KVVTCLLIQSLIIECFAIAAYNIYIPVADDFARKITEGVVKEEYSHLNFGEVWLKEHF<br>AESKAELELANRQNLPIVWKMLNQVEGDAHTMAMEKDALVEDFMIQYGEALS<br>NIGFSTRDIMRLSAYGLIGA* |
| Ado$_{Pm_A134F}$ | MPTLEMPVAAVLDSTVGSSEALPDFTSDRYKDAYSRINAIVIEGEQEAHDNYIAIG<br>TLLPDHVEELKRLAKMEMRHKKGFTACGKNLGVEADMDFAREFFAPLRDNFQT<br>ALGQGKTPTCLLIQALLIEAFAISFYHTYIPVSDPFARKITEGVVKDEYTHLNYGEA<br>WLKANLESCREELLEANRENLPLIRRMLDQVAGDAAVLQMDKEDLIEDFLIAYQ<br>ESLTEIGFNTREITRMAAAALVS* |
| *GmLOX1* | MFSAGHKIKGTVVLMPKNELEVNPDGSAVDNLNAFLGRSVSLQLISATKADAHG<br>KGKVGKDTFLEGINTSLPTLGAGESAFNIHFEWDGSMGIPGAFYIKNYMQVEFFL<br>KSLTLEAISNQGTIRFVCNSWVYNTKLYKSVRIFFANHTYVPSETPAPLVSYREEEL<br>KSLRGNGTGERKEYDRIYDYDVYNDLGNPDKSEKLARPVLGGSSTFPYPRRGRTG<br>RGPTVTDPNTEKQGEVFYVPRDENLGHLKSKDALEIGTKSLSQIVQPAFESAFDLK<br>STPIEFHSFQDVHDLYEGGIKLPRDVISTIIPLPVIKELYRTDGQHILKFPQPHVVQV<br>SQSAWMTDEEFAREMIAGVNPCVIRGLEEFPPKSNLDPAIYGDQSSKITADSLDL<br>DGYTMDEALGSRRLFMLDYHDIFMPYVRQINQLNSAKTYATRTILFLREDGTLKP<br>VAIELSLPHSAGDLSAAVSQVVLPAKEGVESTIWLLAKAYVIVNDSCYHQLMSH<br>WLNTHAAMEPFVIATHRHLSVLHPIYKLLTPHYRNNMNINALARQSLINANGIIE<br>TTFLPSKYSVEMSSAVYKNWVFTDQALPADLIKRGVAIKDPSTPHGVRLLIEDYPY<br>AADGLEIWAAIKTWVQEYVPLYYARDDDVKNDSELQHWWKEAVEKGHGDLK<br>DKPWWPKLQTLEDLVEVCLIIIWIASALHAAVNFGQYPYGGLIMNRPTASRRLLP<br>EKGTPEYEEMINNHEKAYLRTITSKLPTLISLSVIEILSTHASDEVYLGQRDNPHWT<br>SDSKALQAFQKFGNKLKEIEEKLVRRNNDPSLQGNRLGPVQLPYTLLYPSSEEGLT<br>FRGIPNSISI* |
| UndB | MSPSPASLNDQQRAAHIREQVMAHGNALRQRYPILQHQDALGAGILAFALCG<br>MIGSAALYIGGHLPWWACLLLNAFFASLTHELEHDLIHSMYFRKQPLPHNLMLA<br>LVWLARPSTINPWVRRHLHLNHHKVSGSEADMEERAITNGEPWGIARLLMVG<br>DNMMSSFIRWLRAKNPEHRRLILTRTLKVYAPLGLLNWATWYLFLGFHLLDWA<br>AAALGAPIAWSASTLSVMQVVNVAVVVLVGPNVLRTFCLHFVSSNMHYYGDVE |

| | |
|---|---|
| | PGNVIQQTQVLNPWWLWPLQAFCFNFGSSHAIHHFVVKEPFYIRQLTVPFAHR VMREMGVRFNDFGTFARANRWTRRARTQQERASTA* |
| ‘*Ca*FatB3.5 | MHDRKSKRPSMLMDSFGLESIVQEGLEFRQSFSIRSYEIGTDRTASMETLMNYL QETSLNHCKSTGILLDGFGRTPEMCKRDLIWVVTKMKIKVNRYPAWGDTVEINT WFSRLGKIGKGRDWLISDCNTGEILIRATSAYATMNQKTRRLSKLPYEVHQEIAPL FVDSPPVIEDNDLKLHKFEVKTGDSIHKGLTPGWNDLDVNQHVSNVKYIGWILE SMPTEVLETQELCSLALEYRRECGRDSVLESVTAMDPTKVGGRSQYQHLLRLED GTDIVKCRTEWRPKNPGANGAISTGKTSNGNSVS* |
| ‘*Cho*FatB2.2 | MLPDWSRLLTAITTVFVKSKRPDMHDRKSKRPDMLVDSFGLESTVQDGLVFRQ SFSIRSYEIGTDRTASIETLMNHLQETSLNHCKSTGILLDGFGRTLEMCKRDLIWV VIKMQIKVNRYPAWGDTVEINTRFSRLGKIGMGRDWLISDCNTGEILVRATSAY AMMNQKTRRLSKLPYEVHQEIVPLFVDSPVIEDSDLKVHKFKVKTGDSIQKGLTP GWNDLDVNQHVSNVKYIGWILESMPTEVLETQELCSLALEYRRECGRDSVLESV TAMDPSKVGVRSQYQHLLRLEDGTAIVNGATEWRPKNAGANGAISTGKTSNG NSVS* |
| ‘*Uc*FatB1 | MLEWKPKPKLPQLLDDHFGLHGLVFRRTFAIRSYEVGPDRSTSILAVMNHMQE ATLNHAKSVGILGDGFGTTLEMSKRDLMWVVRRTHVAVERYPTWGDTVEVEC WIGASGNNGMRRDFLVRDCKTGEILTRCTSLSVLMNTRTRRLSTIPDEVRGEIGP AFIDNVAVKDDEIKKLQKLNDSTADYIQGGLTPRWNDLDVNQHVNNLKYVAW VFETVPDSIFESHHISSFTLEYRRECTRDSVLRSLTTVSGGSSEAGLVCDHLLQLEG GSEVLRARTEWRPKLTDSFRGISVIPAEPRV* |
| ‘*Cp*FatB1.4 | MFDRKSKRPSMLMDSFGLERVVQDGLVFRQSFSIRSYEICADRTASMETVMNH VQETSLNQCKSIGLLDDGFGRSPEMCKRDLIWVVTRMKIMVNRYPTWGDTIEV STWLSQSGKIGMGRDWLISDCNTGEILVRATSVYAMMNQKTRRFSKLPHEVRQ EFAPHFLDSPPAIEDNDGKLQKFDVKTGDSIRKGLTPGWYDLDVNQHVSNVKYI GWILESMPTEVLETQELCSLTLEYRRECGRDSVLESVTSMDPSKVGDRFQYRHLL RLEDGADIMKGRTEWRPKNAGTNGAISTGKT* |
| FAP | MASAVEDIRKVLSDSSSPVAGQKYDYILVGGGTAACVLANRLSADGSKRVLVLEA GPDNTSRDVKIPAAITRLFRSPLDWNLFSELQEQLAERQIYMARGRLLGGSSATN ATLYHRGAAGDYDAWGVEGWSSEDVLSWFVQAETNADFGPGAYHGSGGPM RVENPRYTNKQLHTAFFKAAEEVGLTPNSDFNDWSHDHAGYGTFQVMQDKGT RADMYRQYLKPVLGRRNLQVLTGAAVTKVNIDQAAGKAQALGVEFSTDGPTGE RLSAELAPGGEVIMCAGAVHTPFLLKHSGVGPSAELKEFGIPVVSNLAGVGQNL QDQPACLTAAPVKEKYDGIAISDHIYNEKGQIRKRAIASYLLGGRGGLTSTGCDR GAFVRTAGQALPDLQVRFVPGMALDPDGVSTYVRFAKFQSQGLKWPSGITMQ LIACRPQSTGSVGLKSADPFAPPKLSPGYLTDKDGADLATLRKGIHWARDVARSS ALSEYLDGELFPGSGVVSDDQIDEYIRRSIHSSNAITGTCKMGNAGDSSSVVDNQ |

| | |
|---|---|
| | LRVHGVEGLRVVDASVVPKIPGGQTGAPVVMIAERAAALLTGKATIGASAAAPA TVAA* |

### Supplementary Table S2. List of strains and plasmids used in this study

| Strain name | Description |
|---|---|
| Pro 1 | ΔphaEC::P$_{trc}$ pduP P$_{trc}$ phaJ ter SpR, ΔphaA::CmR P$_{sca6\text{-}2}$ nphT7 xfpk |
| Pro 2 | ΔphaEC::P$_{trc}$ pduP P$_{trc}$ phaJ ter SpR, ΔphaA::CmR P$_{sca6\text{-}2}$ nphT7 xfpk, ΔNSII::P$_{L22}$ dCas9 GmR P$_{L31}$ [sgRNA slr1192] |
| Pro 3 | ΔphaEC::P$_{trc}$ pduP P$_{trc}$ phaJ ter SpR, ΔphaA::CmR P$_{sca6\text{-}2}$ nphT7 xfpk, ΔNSII::P$_{L31}$ dCas9 GmR P$_{L31}$ [sgRNA slr0942_sll0990_slr0091_slr1192] |
| Pro 1 + Ado$_{Pm}$ | ΔphaEC::P$_{trc}$ pduP P$_{trc}$ phaJ ter SpR, ΔphaA::CmR P$_{sca6\text{-}2}$ nphT7 xfpk, pRSF1010-Ery-Ptrc-ado$_{Pm}$ |
| Pro 2 + Ado$_{Pm}$ | ΔphaEC::P$_{trc}$ pduP P$_{trc}$ phaJ ter SpR, ΔphaA::CmR P$_{sca6\text{-}2}$ nphT7 xfpk, ΔNSII::P$_{L22}$ dCas9 GmR P$_{L31}$ [sgRNA slr1192], pRSF1010-Ery-Ptrc-ado$_{Pm}$ |
| Pro 3 + Ado$_{Pm}$ | ΔphaEC::P$_{trc}$ pduP P$_{trc}$ phaJ ter SpR, ΔphaA::CmR P$_{sca6\text{-}2}$ nphT7 xfpk, ΔNSII::P$_{L31}$ dCas9 GmR P$_{L31}$ [sgRNA slr0942_sll0990_slr0091_slr1192] , pRSF1010-Ery-Ptrc-ado$_{Pm}$ |
| Pro 3 + Ado$_{Pm_A134F}$ | ΔphaEC::P$_{trc}$ pduP P$_{trc}$ phaJ ter SpR, ΔphaA::CmR P$_{sca6\text{-}2}$ nphT7 xfpk, ΔNSII::P$_{L31}$ dCas9 GmR P$_{L31}$ [sgRNA slr0942_sll0990_slr0091_slr1192] , pRSF1010-Ery-Ptrc-ado$_{Pm_A134F}$ |
| Pro 3 + Ado$_{Np}$ | ΔphaEC::P$_{trc}$ pduP P$_{trc}$ phaJ ter SpR, ΔphaA::CmR P$_{sca6\text{-}2}$ nphT7 xfpk, ΔNSII::P$_{L31}$ dCas9 GmR P$_{L31}$ [sgRNA slr0942_sll0990_slr0091_slr1192] , pRSF1010-Ery-Ptrc-ado$_{Np}$ |
| Pro 3 + Ado$_{Pm_2xRBS}$ | ΔphaEC::P$_{trc}$ pduP P$_{trc}$ phaJ ter SpR, ΔphaA::CmR P$_{sca6\text{-}2}$ nphT7 xfpk, ΔNSII::P$_{L31}$ dCas9 GmR P$_{L31}$ [sgRNA slr0942_sll0990_slr0091_slr1192] , pRSF1010-Ery-Ptrc-ado$_{Pm_2xRBS}$ |
| Δ*aas* | Δ*aas*::KmR |
| Δ*aas*-*Gm*LOX1$_{RBS3}$ | Δ*aas*::KmR, pIY1122 pRSF1010-Ery-Ptrc-*Gm*LOX1 (RBS3) |
| Δ*aas*-*Gm*LOX1$_{RBS5}$ | Δ*aas*::KmR, pIY1123 pRSF1010-Ery-Ptrc-*Gm*LOX1 (RBS5) |
| Δ*aas*-P$_{coa}$-'*Ca*FatB3.5 | Δ*aas*::KmR, pIY848 pRSF1010-Ery-Pcoa-'*Ca*FatB3.5 |
| Δ*aas*-P$_{coa}$-'*Ca*FatB3.5-UndB | Δ*aas*::KmR, pIY1197 pRSF1010-Ery-Pcoa-'*Ca*FatB3.5-UndB |
| Δ*aas*-P$_{coa}$-'*Cho*FatB2.2 | Δ*aas*::KmR, pIY783 pRSF1010-Ery-Pcoa-'*Cho*FatB2.2 |
| Δ*aas*-P$_{coa}$-'*Cho*FatB2.2-UndB | Δ*aas*::KmR, pIY1198 pRSF1010-Ery-Pcoa-'*Cho*FatB2.2-UndB |
| Δ*aas*-P$_{coa}$-'*Uc*FatB1 | Δ*aas*::KmR, pIY786 pRSF1010-Ery-Pcoa-'*Uc*FatB1 |
| Δ*aas*-P$_{coa}$-**4**-'*Uc*FatB1-**4**-UndB | Δ*aas*::KmR, pIY868 pRSF1010-Ery-Pcoa-**4**-'*Uc*FatB1-**4**-UndB |
| Δ*aas*-P$_{coa}$-**3**-UndB-**4**-'*Uc*FatB1-4 | Δ*aas*::KmR, pIY1077 pRSF1010-Ery-Pcoa-**3**-UndB-**4**-'*Uc*FatB1-4 |
| Δ*aas*-P$_{coa}$-**4**-'*Uc*FatB1-**3**-UndB | Δ*aas*::KmR, pIY1076 pRSF1010-Ery-Pcoa-**4**-'*Uc*FatB1-**3**-UndB |
| Δ*aas*-P$_{trc}$-**3**-UndB-**4**-'*Uc*FatB1-4 | Δ*aas*::KmR, pIY1078 pRSF1010-Ery-Ptrc-**3**-UndB-**4**-'*Uc*FatB1-4 |
| Δ*aas*-P$_{coa}$-**5**-'*Uc*FatB1-**5**-UndB | Δ*aas*::KmR, pIY869 pRSF1010-Ery-Pcoa-**5**-'*Uc*FatB1-**5**-UndB |
| Δ*aas*-P$_{coa}$-**3**-'*Uc*FatB1-**3**-UndB | Δ*aas*::KmR, pIY867 pRSF1010-Ery-Pcoa-**3**-'*Uc*FatB1-**3**-UndB |
| Δ*aas*-P$_{coa}$-**4**-'*Cp*FatB1.4-Ptrc-**3**-FAP | Δ*aas*::KmR, pIY1001 pRSF1010-Ery-Pcoa-**4**-'*Cp*FatB1.4-Ptrc-**3**-FAP |
| Δ*aas*-P$_{trc}$-**4**-FAP | Δ*aas*::KmR, pIY646 pRSF1010-Ery-Ptrc-**4**-FAP |
| Δ*aas*-P$_{trc}$-**1**-FAP-**4**-'*Uc*FatB1 | Δ*aas*::KmR, pIY1079 pRSF1010-Ery-Ptrc-**1**-FAP-**4**-'*Uc*FatB1 |

| | |
|---|---|
| Δ*aas*-$P_{trc}$-**2**-FAP-**4**-'*Uc*FatB1 | Δ*aas*::KmR, pIY1080 pRSF1010-Ery-Ptrc-**2**-FAP-**4**-'*Uc*FatB1 |
| Δ*aas*-$P_{trc}$-**3**-FAP-**4**-'*Uc*FatB1 | Δ*aas*::KmR, pIY1081 pRSF1010-Ery-Ptrc-**3**-FAP-**4**-'*Uc*FatB1 |
| Δ*aas*-$P_{trc}$-**4**-FAP-**4**-'*Uc*FatB1 | Δ*aas*::KmR, pIY1082 pRSF1010-Ery-Ptrc-**4**-FAP-**4**-'*Uc*FatB1 |
| Δ*aas*-$P_{trc}$-**5**-FAP-**4**-'*Uc*FatB1 | Δ*aas*::KmR, pIY1083 pRSF1010-Ery-Ptrc-**5**-FAP-**4**-'*Uc*FatB1 |
| Δ*aas*-$P_{trc}$-**6**-FAP-**4**-'*Uc*FatB1 | Δ*aas*::KmR, pIY1084 pRSF1010-Ery-Ptrc-**6**-FAP-**4**-'*Uc*FatB1 |

Number in bold represents a ribosome binding site.

RBS sequence used in this study

| RBS No. | RBS sequence | Source |
|---|---|---|
| 1 | ATCACACAGGAC | BBa_B0033[$] |
| 2 | AAAGAGGGGAAA | BBa_B0064[$] |
| 3 | AAAGAGGAGAAA | BBa_B0034[$] |
| 4 | ATCACAAGGAGG | Shine-Dalgarno (SD) *E. coli* consensus |
| 5 | ATTAGTGGAGGT | Anti-SD complementary sequence (RBS*)[§] |
